# Stimulus-reward contingencies drive long-lasting alterations in neocortical somatostatin inhibition during learning

**DOI:** 10.1101/2023.12.08.570881

**Authors:** Eunsol Park, Dika A. Kuljis, Mo Zhu, Joseph A. Christian, Alison L. Barth

**Affiliations:** Department of Biological Sciences, Carnegie Mellon University, 4400 Fifth Ave., Pittsburgh PA 15232; Biology Department, Sacred Heart University, 5151 Park Ave., Fairfield, CT 06825

## Abstract

Learning involves the association of discrete events in the world to infer causality, likely through a cascade of changes at input- and target-specific synapses. Transient or sustained disinhibition may initiate cortical circuit plasticity important for association learning, but the cellular networks involved have not been well-defined. Here we show that sensory association learning drives a durable, target-specific reduction in inhibition from somatostatin (SST)-expressing GABAergic neurons onto pyramidal (Pyr) neurons in superficial but not deep layers of mouse somatosensory cortex. Critically, SST-output was not altered when stimulus and rewards were unpaired, indicating that these neurons are not modified by sensory input alone. Depression of SST output onto Pyr neurons could be phenocopied by chemogenetic suppression of SST activity outside of the training context. Thus, neocortical SST neuron output is persistently modified by convergent sensory and reinforcement signals to selectively disinhibit superficial layers of sensory neocortex during learning.

## Introduction

Disinhibition is thought to facilitate network rewiring during learning (Barron, 2021; Letzkus et al., 2015; Makino and Komiyama, 2015). Although brain states, such as task engagement or reinforcement cues, can dynamically suppress inhibition (Fu et al., 2014; Kato et al., 2015; Kuchibhotla et al., 2017; Lee et al., 2013; Pi et al., 2013), long-lasting changes in inhibitory synaptic output may also play a role (Chen et al., 2015; Kida et al., 2016; Makino and Komiyama, 2015; Sarro et al., 2015; Schroeder et al., 2023; Yang et al., 2022). Inhibitory synaptic plasticity can thus provide a potent means to reduce inhibitory output, and *in vitro* studies have established specific requirements for the durable alteration in inhibitory synaptic strength (Chiu et al., 2019; McFarlan et al., 2023). For example, neocortical somatostatin-expressing (SST) inputs to pyramidal (Pyr) neurons can be potentiated by the activation of NMDA-type glutamate receptors, and these synapses can undergo spike-timing-dependent plasticity (Chiu et al., 2013; Udakis et al., 2020). Despite studies that establish specific requirements for inhibitory synapse plasticity *in vitro*, a connection between these fine-scale phenomena and pathway-specific disinhibition during a natural learning behavior has not been well-established.

*In vivo* monitoring of inhibitory neuron firing has revealed changes in activity during sensory and motor learning (Cummings et al., 2022; Makino and Komiyama, 2015; Ren et al., 2022). Cell-type specific analysis of inhibitory neuron firing indicates that both SST and parvalbumin-expressing (PV) interneurons can become more selective during sensory learning (Khan et al., 2018), and that sensory association learning and passive sensory exposure differentially affect SST and PV neurons (Kato et al., 2015; Makino and Komiyama, 2015). Thus, plasticity of inhibition is sensitive to both cell identity and also training conditions.

*In vivo* imaging and recording studies indicate that SST neurons are distinctively engaged during sensory perception and learning (Gentet et al., 2012; Sachidhanandam et al., 2016; Yu et al., 2019). Prior studies suggest that stimulus-evoked SST activity may be reducing during learning (Makino and Komiyama, 2015; Yang et al., 2022), an observation consistent with dynamic regulation of SST firing during stimulus-reward pairing. A reduction in interneuron activity during learning implicates upstream inhibitory networks that can control SST activity (Jiang et al., 2015; Kepecs and Fishell, 2014; Pfeffer et al., 2013). For example, synaptic inhibition from vasoactive-intestinal peptide (VIP)-expressing GABAergic neurons onto SST neurons is a common circuit observed across both sensory and motor cortex, and both VIP and SST neurons are sensitive to neuromodulators that are released during reinforcement learning (Hangya et al., 2015; Kuchibhotla et al., 2017; Muñoz et al., 2017; Urban-Ciecko et al., 2018). However, whether dynamic changes in SST neuron activity during behavior are sufficient to facilitate a durable restructuring of inhibition, as well as the layer- and target-cell specificity of this plasticity, has not been investigated. Therefore, we sought to examine whether learning of a sensory association task could drive SST-mediated inhibitory synaptic plasticity in neocortical circuits.

We took advantage of a high-throughput training environment for whisker-dependent sensory association learning (Bernhard et al., 2020) to investigate how SST synaptic output in primary somatosensory (barrel) cortex might be altered during acquisition of the task. Using SST-targeted ChR2 expression, we quantified the magnitude of SST output for Pyr neurons in both L2 and L5, two cortical layers where sensory learning-related changes have been well-documented in prior studies (Audette et al., 2019; Doron et al., 2020; Khan et al., 2018; Kuhlman et al., 2014; Lacefield et al., 2019). We found that SST output on L2 Pyr neurons was rapidly reduced at the onset of sensory association training, and that these changes were selectively induced by stimulus-reward pairing but not during pseudotraining where stimulus and rewards were unpaired. Chemogenetic suppression of SST activity was sufficient to reduce SST output compared to control animals, consistent with a learning-related reduction in stimulus-evoked SST activity (Makino and Komiyama, 2015; Yang et al., 2022). Furthermore, the high-throughput training paradigm enabled us to detect behavioral correlations of SST disinhibition at different stages of learning across a large cohort of animals, where reduced SST output and stimulus-reward association became strongly correlated after several days of training. We propose that reduced SST inhibition onto L2 neurons in primary sensory cortex is a critical early step in neocortical circuit plasticity that is linked to learning stimulus-reward associations.

## Results

### Automated training drives association learning

To assess changes in inhibitory synaptic strength, we took advantage of a high-throughput, homecage behavioral training paradigm for sensory association learning (Bernhard et al., 2020). SST-Cre transgenic mice were acclimated to the training cage without a sensory stimulus for 1-2 days (Figure 1A, B). At the onset of SAT, a gentle (∼6 psi, 500ms airpuff) was delivered when animals positioned themselves at the nosepoke prior to water delivery (Figure 1B). For 20% of initiated trials, neither an airpuff nor a water reward was delivered (“blank” trials). To assess whether the animals had learned that the airpuff predicted the subsequent water delivery, we compared anticipatory licking between the end of the airpuff and the delivery of water in stimulus-reward versus “blank” trials.

**Figure 1.**
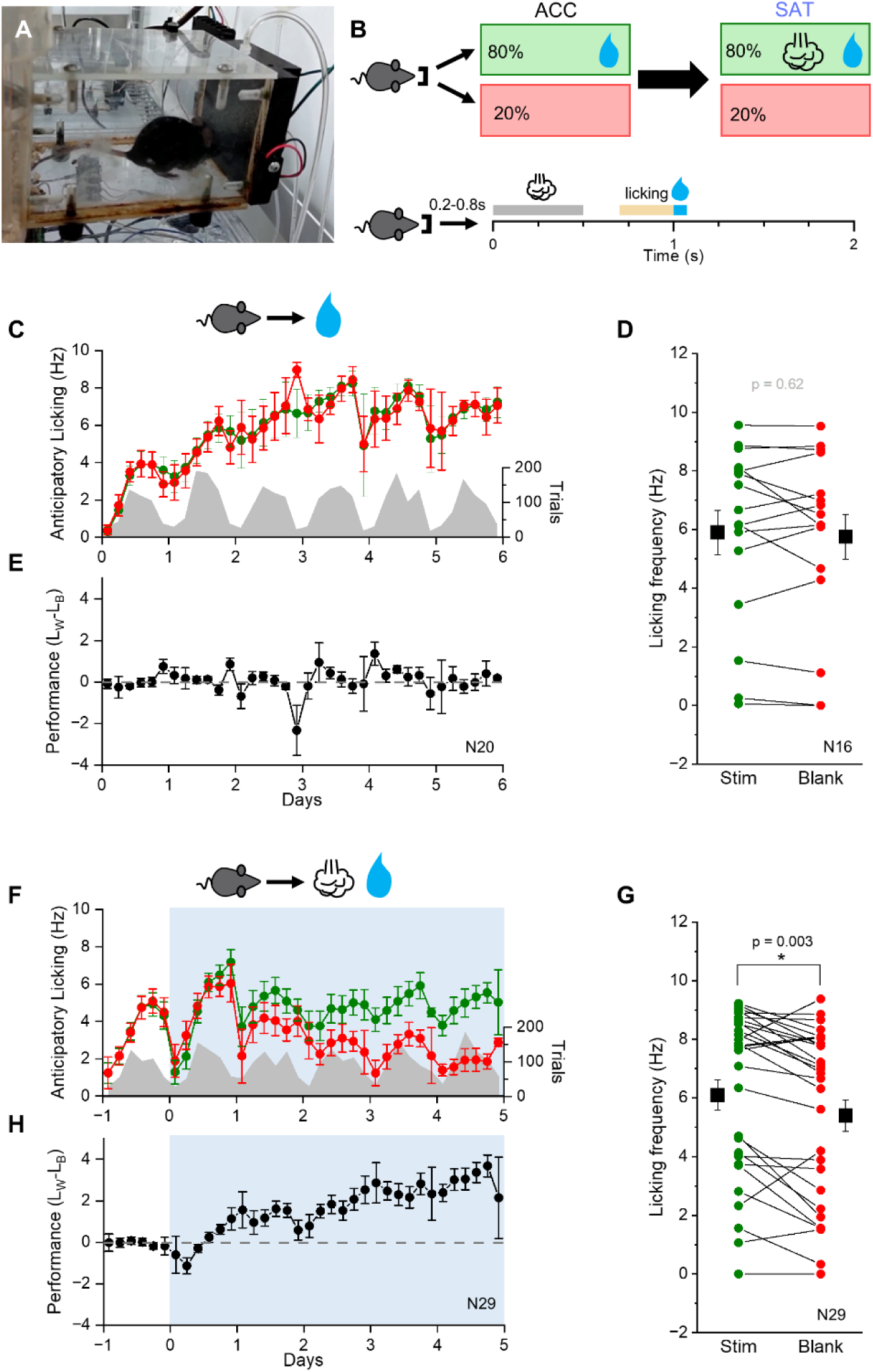
Mice learn to associate whisker stimuli with water after sensory association training. (A) Automated cage training set-up. (B) Top: timeline of cage acclimation and training. Sensory association training (SAT) starts after 1-2 days of acclimation period. Bottom: trial structure. Nosepoke initiates trial onset, where a random delay time (0.2-0.8s) is followed by a gentle airpuff (500ms, 6psi), a fixed 500ms delay, and then water delivery. Anticipatory licking frequency (Hz) 300ms before water delivery (yellow line) was calculated for both stimulus and blank trials to evaluate animals’ behavior. (C) Mean anticipatory licking frequency (Hz) stimulus absent, water trials (green) and blank trials (red) after 1-6 days of acclimation (ACC). Blue shade indicates the training period. The distribution of trial numbers over days is shown in gray. 1 day of ACC, N=20 mice; 2 days of ACC N=16 mice; 6 days of ACC N=6 mice. (D) Mean anticipatory licking frequency (Hz) for the last 20% of stimulus (green) and blank (red) trials for each animal after 2 days of ACC. Group mean±SEM shown in black symbols (Stim: 5.9±0.8Hz vs. Blank: 5.7±0.8Hz; N=16 mice). (E) Performance (licking_stim/water_ - licking_blank_; see methods) was calculated for each animal and averaged across all 4-hr time bins. (F-H) As in (C-E), but for animals after sensory association training (SAT). 1 day of SAT (SAT1), N=29 mice; 5 days of SAT (SAT5), N=10 mice. (G) Group mean±SEM shown in black symbols (stim: 6.3±0.5Hz vs. blank: 5.6±0.5Hz; N=29 mice). Lick frequencies were compared using a paired sample Wilcoxon signed rank test. *P–*value in figure. \**p*<0.05.

Over the course of several days, animals learn to associate a whisker stimulus with a water reward, measured by an increase in lick frequency to stimulus compared to “blank” trials (Figure 1F). Prior studies have shown that this sensory association training (SAT) is whisker-dependent (Bernhard et al., 2020) and also drives rapid changes in thalamocortical synaptic strength (Audette et al., 2019; Ray et al., 2023), indicating that this training is sufficient to initiate both synaptic and behavioral plasticity. Across the population of trained SST-Cre animals, mice showed a significantly greater lick frequency in stimulus versus “blank” trials after just 1 day of SAT (Figure 1G; 6.3±0.5Hz versus 5.6±0.5Hz). The relative difference in lick frequency could be represented as performance by calculating the difference between stimulus- and “blank”-associated licking (Figure 1H; performance=licking_stim/water_ - licking_blank_; see methods). Animals housed in the training environment but without a predictive cue to indicate water delivery showed no difference in anticipatory licking between water and “blank” trials (Figure 1A-C).

### SST-mediated inhibition onto L2 Pyr neurons is reduced during SAT

To assess the strength of SST-mediated synaptic inhibition in barrel cortex, we prepared acute brain slices from SST-Cre x Ai32 (flex-ChR2) mice that had undergone cage acclimation (ACC) only or SAT after acclimation (Figure 2A). Pyr neurons from superficial and deep layers were targeted for whole-cell patch-clamp recordings. SST- inhibitory postsynaptic currents (IPSCs) were evoked using brief pulses of blue light (5 ms, 0.1Hz), and the mean amplitude of the evoked response was compared across superficial and deep layers of neocortex for each condition. Analysis in acute brain slices enabled us to assess whether there were long-lasting changes in SST-mediated input outside of behavior, as well as to examine the strength of SST output in deep layers, a cortical area that has not been well-investigated *in vivo*.

**Figure 2.**
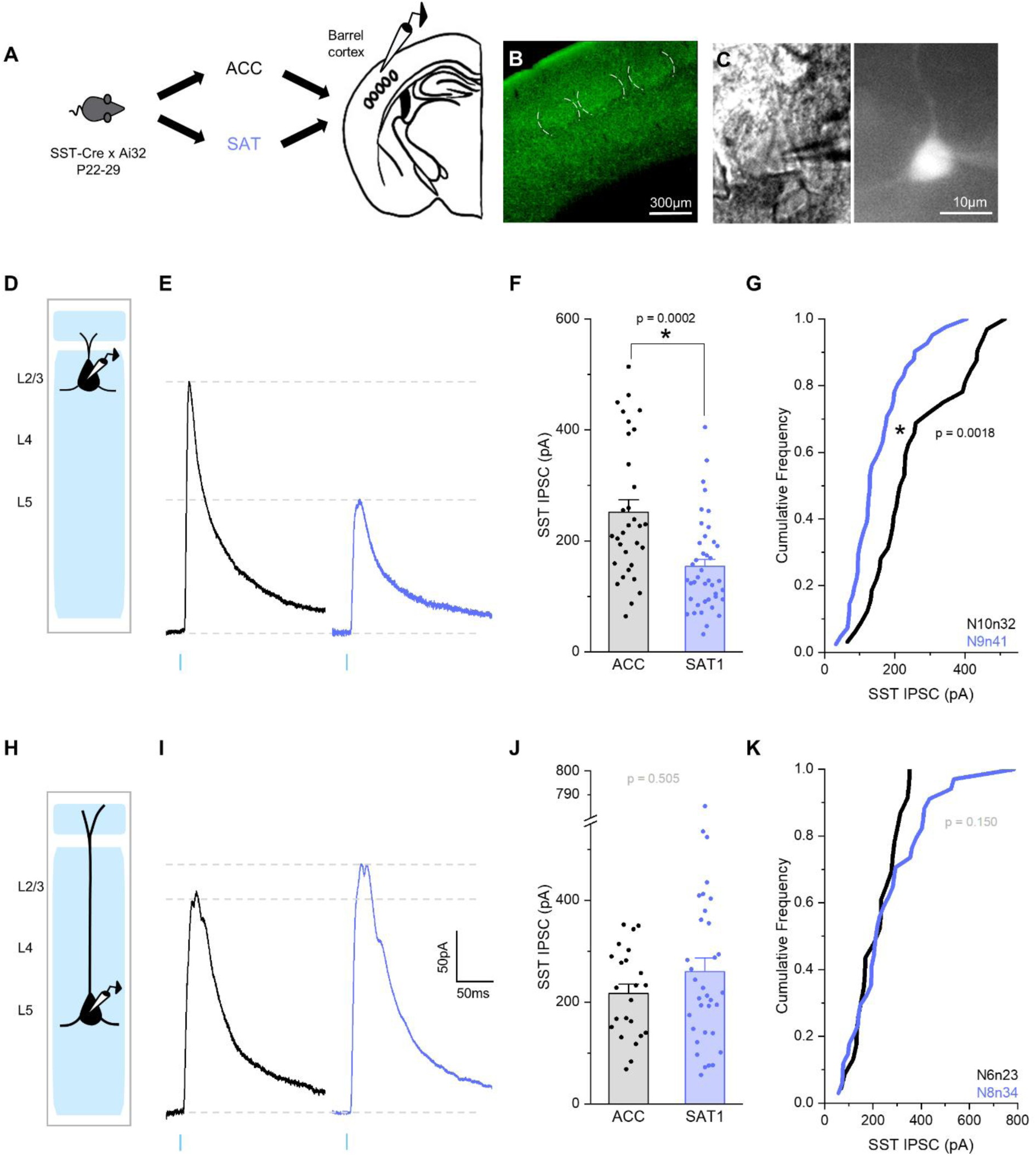
SST inhibition on L2 Pyr neurons is reduced after a single day of training. (A) Schematic of training conditions and tissue preparation for SST-Cre x Ai32 mice. (B) An example image of the barrel cortex in an acute brain slice, where SST neurons in all layers are expressing ChR2. White dashed lines indicate outlines of the barrels. (C) Pyr neuron identity was verified by regular-spiking firing phenotype and morphology (left: a brightfield image; right: a Pyr neuron with cell fill after recording). (D) Schematic of recording from L2 Pyr neurons. (E) Example ChR2-evoked SST-IPSC (10 sweep average) recorded from a L2 Pyr neuron after 1-2 days of acclimation (ACC; black) and 1 day of SAT (SAT1; blue). (F) Peak amplitude of SST-IPSCs for L2/3 Pyr neurons after ACC (252.2±21.9pA; N=10 mice, n=32 cells) and SAT1 (154.1±13.0pA; N=9 mice, n=41 cells). Bar graphs represent mean+SEM. Mann-Whitney U test (U=987, n=32 and 41, two-tailed). (G) Cumulative distribution histogram of SST-IPSC amplitudes for L2 Pyr neurons from ACC and SAT1 mice. Komolgorov-Smirnov (K-S) test. (H-K) Same as (D)-(G), but for L5 Pyr neurons (ACC: 216.8±18.1pA; N=6 mice, n=23 cells vs. SAT1: 250.4±27.4pA; N=8 mice, n=34 cells). Mann-Whitney U test (U=349.5, n=23 and 34, two-tailed). K-S test. *P–*value in figure. \**p*<0.05.

After just 1 day of SAT, the mean amplitude of the SST-IPSC onto L2 Pyr neurons was reduced to ∼40% of control values, a difference that was highly significant (Figure 2F; ACC 252.2±21.9pA versus SAT1 154.1±13.0pA). This was not the case for L5 Pyr neurons, where the SST-IPSC was slightly larger after training, a difference that was not significant (Figure 2J; ACC 216.8±18.1pA versus SAT1 259.4±27.4pA). SST output to PV inhibitory neurons in superficial layers was not reduced by SAT, indicating target-specific effects (Figure S1; ACC 119.9±22.5pA versus SAT1 105.5±14.8pA).

The reduction in SST-IPSC amplitude after training could not be attributed to reduced intrinsic excitability of L2 SST neurons, since SST neurons were marginally more excitable after SAT, particularly at the highest levels of current injection. There were no significant changes in resting membrane potential, input resistance, or rheobase current for SST neurons (Figure 3). The decrease in SST-IPSCs was not sex-dependent, as the magnitude of the reduction was similar across males and females (Figure S2; 40% decrease in both males and females from ACC to SAT1). Thus, these data indicate that there is a rapid and layer-specific reduction in SST output in barrel cortex at the early stages of a tactile learning task that is preserved outside of the task context, in acute brain slices from trained animals.

**Figure 3.**
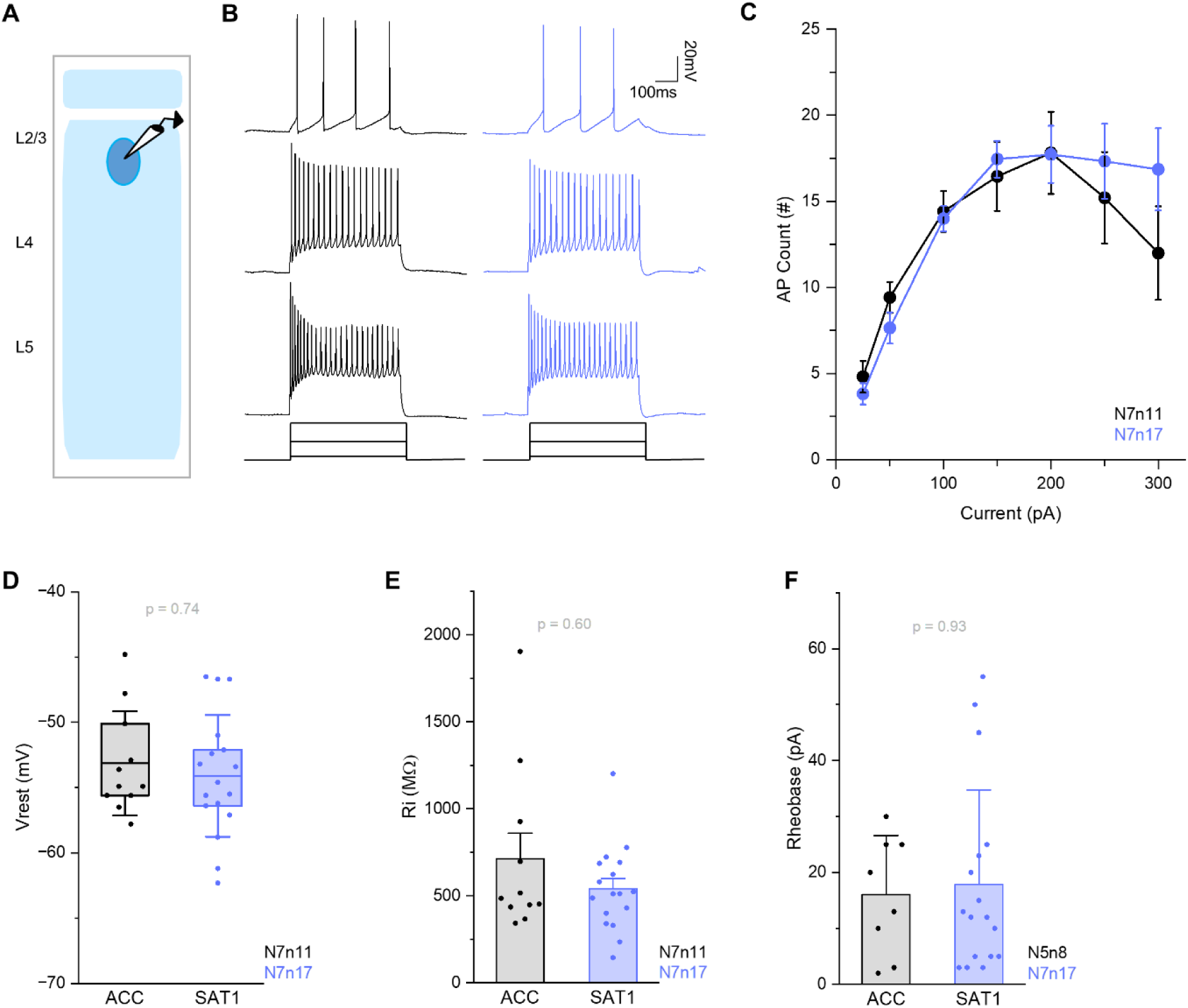
Intrinsic properties of SST are not altered by training. (A) Schematic of the experiment setup. Either eYFP-expressing or ChR2-expressing SST neurons were targeted. (B) Representative firing response following 25pA, 150pA, and 300pA current injection (500ms) recorded from L2/3 low-threshold spiking SST neurons after 1-2 days of acclimation (ACC: black) and 1 day of SAT (SAT1: blue). (C) F-I curve of L2/3 SST neurons in ACC (black) and SAT1 (blue) animals. Mean±SEM. Two-way repeated measures ANOVA F_(1,25)_=0.26, *p*-value=0.6. Tukey’s *p*-value in S1 Table. (D) Resting membrane potential comparison for ACC (−53.1±4.0mV; N=7 mice, n=11 cells) and SAT1(−54.1±4.7mV; N=8 mice, n=17 cells). Box is 25th and 75th quartile, whiskers are SD, and midline is mean. Mann-Whitney U test (U=101, n=11 and 17, two-tailed). (E) Input resistance comparison (mean+SEM) for ACC (713.9±145.9MΩ; N=7 mice, n=11 cells) and SAT1 (541.3±59.1MΩ; N=8 mice, n=17 cells). Mann-Whitney U test (U=105, n=11 and 17, two-tailed). (F) Rheobase comparison (mean+SD) for ACC (16±10.6pA, N=5 mice, n=8 cells) and SAT1 (17.9±16.9pA, N=8 mice, n=17 cells). Rheobase was recorded from 8 out of 11 cells from ACC animals. Mann-Whitney U test (U=70, n=8 and 11, two-tailed). *P–*value in figure.

### Stimulus-reward coupling is required to induce SST-output depression

SAT in our behavioral paradigm pairs the whisker stimulus with a temporally-delayed water reward. To evaluate the effects of repeated stimulation without consistent reward pairing on the SST-IPSC amplitude in L2 Pyr neurons, we designed a pseudotraining (PSE) protocol where the same fraction of trials (80%) included the whisker stimulus, but the reward was delivered on a separate schedule (Audette et al., 2019). Thus, animals received an identical fraction of stimulus and reward trials but without a consistent association of the stimulus with the reward (Figure 4A).

**Figure 4.**
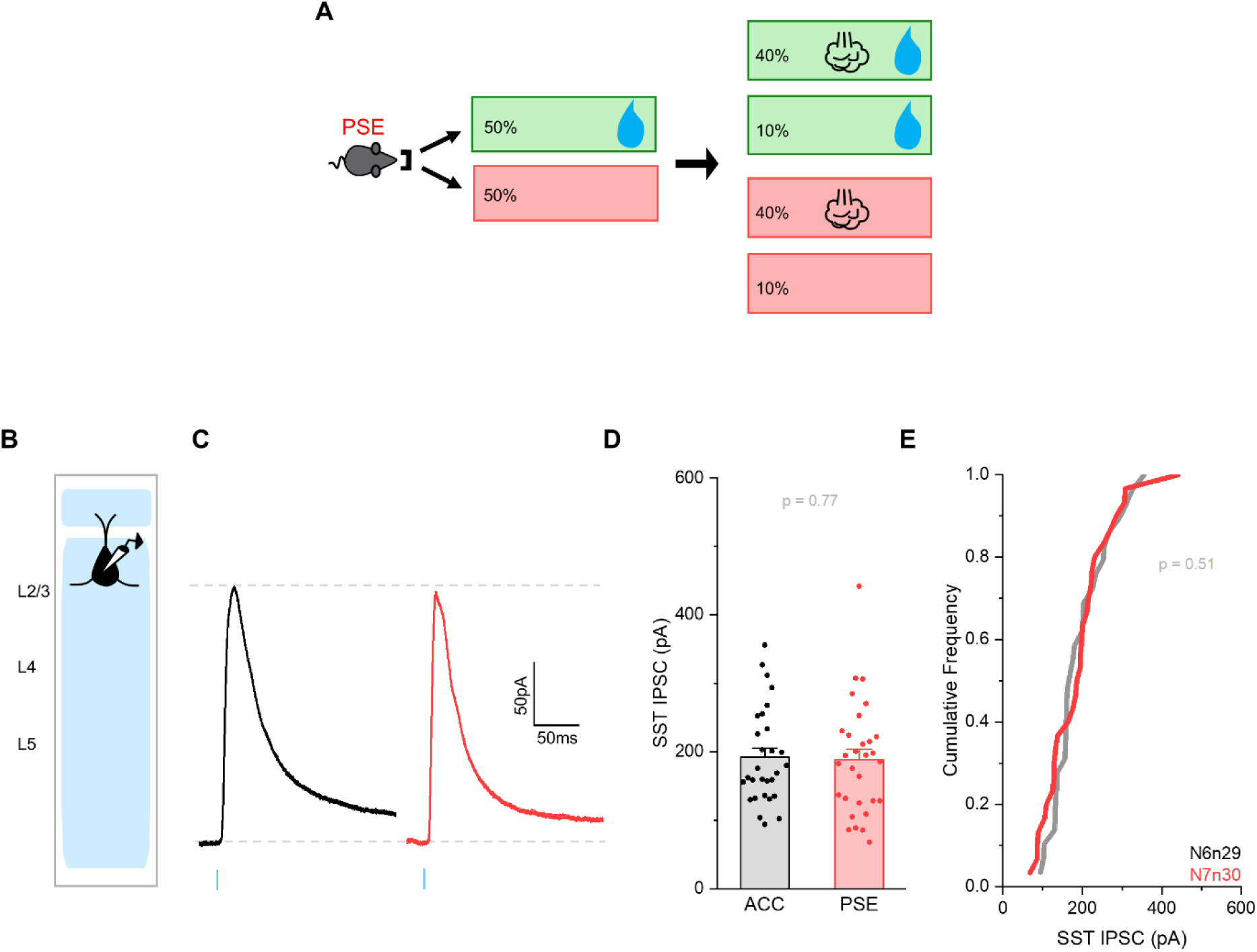
Uncoupled stimulus and reward during pseudotraining does not alter SST inhibition onto L2 Pyr neurons. (A) Schematic of a pseudotraining (PSE) trial structure. Animals were acclimated in the training cage without airpuff for 1 day prior to pseudotraining onset. (B) Schematic of recording from L2 Pyr neurons in SST-Cre x Ai32 mice. (C) Example ChR2-evoked SST-IPSC (10 sweep average) recorded from a L2 Pyr neuron after 1-2 days of acclimation (ACC; black) and 1 day of pseudotraining (PSE; red). (D) Peak amplitude of SST-IPSCs for L2 Pyr neurons from ACC (192.1±13.0pA; N=6 mice, n=29 cells) and PSE (188.6±15.0pA; N=7 mice, n=30 cells) mice. Bar graphs represent mean+SEM. Mann-Whitney U test (U=455, n=29 and 30, two-tailed). (E) Cumulative distribution histogram of SST-IPSC amplitudes for L2 Pyr neurons from ACC and PSE mice. K-S test. *P–*value in figure.

Under these conditions, SST output onto L2 Pyr neurons was unchanged compared to mice kept in the training cage but without the sensory stimulus (Figure 4C-E; ACC 192.1±13.0pA versus PSE 188.6±15.0pA). Notably, animals typically received a greater number of stimulus trials during pseudotraining since the fraction of trials with water delivery was only 50% (compared to 80% for SAT), and freely-moving mice were motivated to complete a sufficient number of trials so that daily water consumption was similar across conditions (∼2.5 mL/day). Thus, sensory stimulation is not sufficient to drive a reduction in SST output, and the consistent presentation of sensory stimuli with reward information is required for SST-output depression.

### Chemogenetically reducing SST activity is sufficient to depress SST output

*In vivo* and *in vitro* recordings show that during sensory and thalamocortical stimulation, SST neurons exhibit at least a short-term reduction in firing activity (Audette et al., 2018; Gentet et al., 2012; Yu et al., 2019), consistent with activation of local inhibitory neurons that synapse onto these cells. It is thus reasonable to hypothesize that the activity of SST neurons may be dynamically suppressed during stimulus-reward pairing and that the repeated activation of this pathway could drive long-lasting changes in SST output.

What are the signals during reward-association learning that drive SST output depression? Candidate mechanisms could be neuromodulators that directly act upon SST neurons or synaptic mechanisms, including local interneurons to suppress SST activity, such as VIP neurons or, to a lesser extent, PV and neurogliaform cells. Alternatively, because L2/3 SST neurons are densely innervated by local Pyr neurons (Urban-Ciecko and Barth, 2016), the signal for SST output plasticity might be related to changes in the level or timing of Pyr activity. In all these scenarios, the activity of SST neurons is suppressed by training-related signals.

We took advantage of chemogenetic (Designer Receptors Exclusively Activated by Designer Drugs; DREADDs) approaches to reduce SST activity, well-suited to the freely-moving condition of our training paradigm since the chemogenetic activator Clozapine-N-Oxide (CNO) could be delivered non-invasively via the drinking water. This specific control of SST neurons also obviated questions about direct or indirect regulation of firing during reward-association learning, focusing the experimental question on whether decreasing SST activity was sufficient to drive output depression. To calibrate the effects of CNO on stimulus-evoked SST firing, AAV viruses encoding Cre-dependent hM4D(Gi) were stereotaxically injected into the barrel cortex of SST-Cre x Ai148 (Flex-GCaMP6f) transgenic mice at the same time as cranial windows were implanted for subsequent monitoring of SST Ca^++^ transients. SST activity in awake animals was compared before and after intraperitoneal (*i.p.*) administration of CNO. Within 30 minutes of administration, the airpuff-driven activity of L2 SST neurons was reduced by 50% (Figure S3), an effect that was not observed with saline administration. Additional experiments targeting L2 SST neurons in acute brain slices showed that bath application of CNO significantly suppressed evoked firing in response to current injection, particularly at the highest levels. Thus, hM4D(Gi)-expression in the presence of CNO significantly reduces evoked firing in SST neurons.

Although excitatory DREADDs have been used to increase target cell activity (for example, hM3D(q)), CNO application did not drive any significant increase in evoked firing for SST neurons with hM3D(q) either *in vitro* or *in vivo* (Figure S4), despite a modest depolarization of membrane potential in the presence of CNO in these inhibitory neurons Figure S4G). It is possible that the unique membrane properties of low-threshold firing SST cells in L2 interfere with the membrane depolarization properties of CNO-activated hM3D(q). We conclude that increasing the activity of SST neurons using excitatory DREADDs must be carefully validated for brain area and experimental conditions, but was not possible in our preparation.

Because inhibitory DREADDs effectively suppressed SST activity, we targeted SST neurons in somatosensory cortex for hM4D(Gi) expression and CNO administration to determine whether this could reduce their inhibition onto Pyr neurons. AAV viruses encoding Cre-dependent hM4D(Gi) were injected into the barrel cortex of SST-Cre x Ai32 transgenic mice. For comparison, a separate group of SST-Cre x Ai32 mice was injected with Cre-dependent mCherry. In order to determine how suppression of SST activity would influence the SST-IPSC in neocortical Pyr neurons, injected mice were housed in the training cage without airpuff stimulation. CNO was delivered through drinking water for two days (∼1mg/kg/day), an approach that has been effective in other studies (Manvich et al., 2018). Importantly, CNO in the drinking water did not significantly change the amount of water consumed or the mean number of trials during the cage acclimation period compared to uninjected animals (hM4D(Gi) 344±14 trials and mCherry 327±9 trials versus ACC, 385±25 trials; p≥0.6 for all comparisons by one-way ANOVA; N=5, 7, 7 mice respectively).

Under these conditions, light-evoked SST-IPSCs in L2 Pyr neurons were significantly reduced compared to mCherry controls (Figure 5D-F; hM4D(Gi)+CNO 126.5±14.2pA versus mCherry+CNO 204.6±20.32pA). Thus, reducing SST activity by itself can initiate SST-IPSC depression.

**Figure 5.**
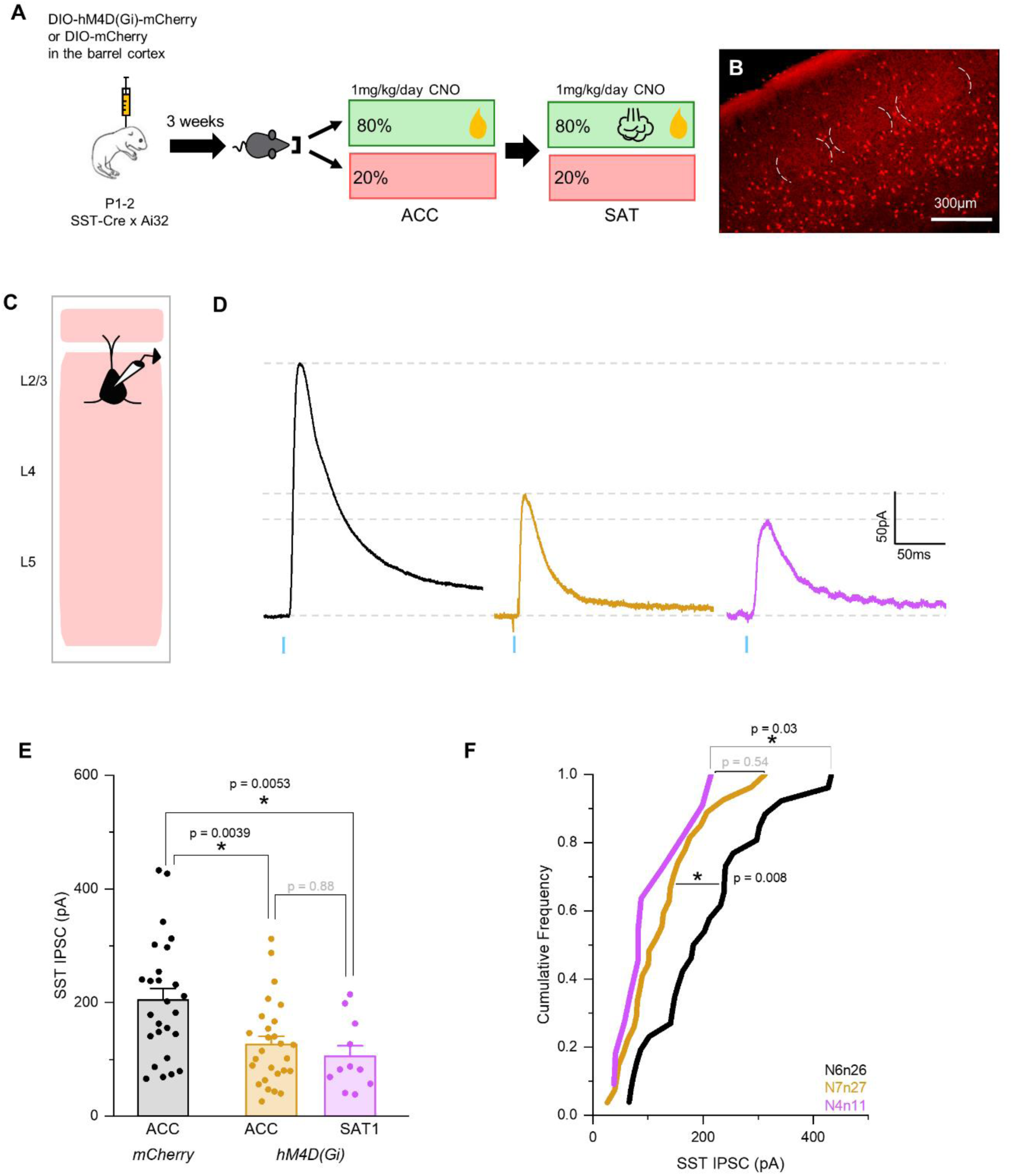
Chemogenetic suppression of SST activity is sufficient to reduce SST inhibition onto L2 Pyr neurons. (A) Schematic of experimental design. Cre-dependent mCherry or hM4D(Gi) virus was injected in the barrel cortex of SST-Cre x Ai32 neonates (P1-2). Mice were acclimated in the training cage, where drinking water contained CNO. A subset of mice was trained in SAT with CNO. (B) An example image of virally-transduced hM4D(Gi)-expression in brain tissue. White dashed lines indicate outlines of the barrels. (C) Schematic of recording from L2 Pyr neurons in mCherry or hM4D(Gi)-expressing SST-Cre x Ai32 mice. (D) Example ChR2-evoked SST-IPSC (10 sweep average) recorded from mCherry-injected mice after 1-2 days of acclimation (mCherry ACC; black), hM4D(Gi)-injected mice after 1-2 days of acclimation (hM4D(Gi) ACC; yellow) and hM4D(Gi)-injected mice after 1 day of SAT (hM4D(Gi) SAT1; purple). All groups were treated with CNO. (D) Peak amplitude of SST-IPSCs for L2 Pyr neurons from mCherry ACC (204.6±20.3pA; N=6 mice, n=26 cells), hM4D(Gi) ACC (126.5±14.2pA; N=7 mice, n=27 cells), and hM4D(Gi) SAT1 (105.4±18.6pA; N=4 mice, n=11 cells) mice. Bar graphs represent mean+SEM. Two-way ANOVA, F_(1,60)_=10.5, *p*=0.002, Tukey’s *p*-value in figure. (E) Cumulative distribution histogram of SST-evoked IPSC amplitudes for all cells from the experimental groups. K-S test. *P*-value in figure. For all panels, \**p*<0.05.

Whisker-association training in these SST-hM4D(Gi) mice, with continued exposure to CNO during both the acclimation period and the onset of SAT, modestly reduced SST-IPSCs in L2 Pyr neurons, a difference that was not significant compared to mean SST-IPSC amplitude for control animals with SST-hM4D(Gi) exposed to CNO (Figure 5D-F; SAT1 hM4D(Gi)+CNO 105.4±18.6pA versus ACC hM4D(Gi)+CNO 126.5±14.2pA). These results indicate that decreasing activity in SST neurons is sufficient to depress SST synaptic output, even in the absence of stimulus-reward coupling. In addition, chemogenetic suppression of SST activity reduced the magnitude of SAT’s effects in depressing SST output compared to control experiments (Figure 2, 39% reduction vs baseline in the SST-IPSC after SAT versus Figure 5, hM4D(Gi)+CNO with a 17% reduction from CNO-treated baseline after SAT). These data suggest that the cellular and synaptic mechanisms involved may be similar between these two conditions.

### Chemogenetic silencing of SST neurons facilitates association learning

SST output depression might be an early and important step that enables both circuit plasticity across the neural circuit linking whisker stimulation and reward and also behavioral change. We hypothesized that the chemogenetic-induced reduction in SST-IPSCs during acclimation to the training environment might facilitate the acquisition or stabilization of a sensory-reward association.

To test this hypothesis, we compared learning trajectories in SST-Cre mice injected with either hM4D(Gi) or mCherry that had been treated with CNO during both the cage acclimation period and then also for the first day of SAT. In contrast to SST-Cre x Ai32 mice (Figure 1), animals with mCherry in SST neurons treated with CNO during both the acclimation and early training period (SAT1) did not exhibit a significant overall difference in anticipatory licking to stimulus versus blank trials at the end of the first day of training (Figure 6A-C; 5.1±0.6Hz versus 4.4±0.5Hz). Indeed, barely half of the trained animals (57%; 8/14 mice) showed greater licking in stimulus versus blank trials, suggesting that CNO treatment itself might influence learning. Although off-target effects of CNO from clozapine metabolism have been described (Jendryka et al., 2019), the dose of CNO used in the current study was calculated to be below the interoceptive threshold for clozapine, a metabolite of CNO (Manvich et al., 2018). Nonetheless, these data suggest that there may be some behavioral effects due to off-target actions of CNO or its metabolites.

**Figure 6.**
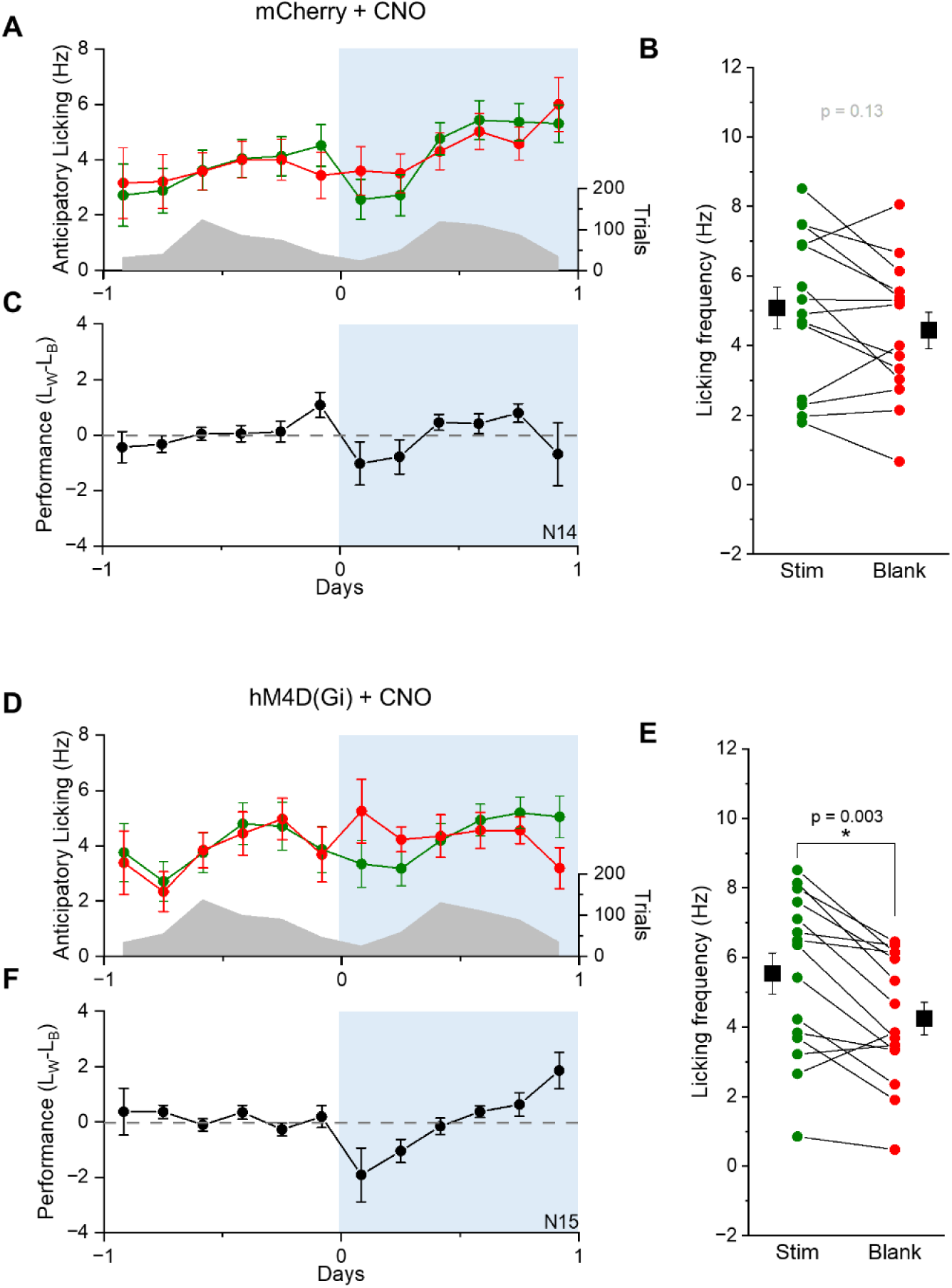
Reduced SST inhibition accelerates learning. (A) A plot of average anticipatory licking frequency (Hz) for stimulus (green) and blank (red) trials for mCherry-injected SST-Cre mice after 1 day of SAT (SAT1). Blue shade indicates the training period. The distribution of average trial numbers over training days is represented as gray shades. (B) The comparison of mean anticipatory licking frequency (Hz) for the last 20% of stimulus (green) and blank (red) trials after 1 day of SAT for mCherry-injected mice. Group mean±SEM shown in black symbols. (Stim: 5.1±0.6Hz vs. Blank: 4.4±0.5Hz; N=14 mice) (C) Average performance for each time-bin (licking_stim/water_ - licking_blank_; see methods). (D-F) As in (A)-(C), but for hM4D(Gi)-injected animals. (Stim: 5.5±0.6Hz vs. Blank: 4.2±0.5Hz; N=15 mice). Lick frequencies were compared using a paired-sample Wilcoxon signed rank test. *P–*value in figure. \**p*<0.05.

Notably, after training mice with hM4D(Gi) in SST neurons and treated with CNO, we observed a significant increase in anticipatory licking to stimulus versus blank trials (Figure 6D-F; 5.5±0.6Hz versus 4.2±0.5Hz). In contrast to the mCherry control animals, almost 90% (13/15) of hM4D(Gi)-SST animals showed an increase in stimulus-evoked anticipatory licking after a single day of training. These data suggest that although CNO treatment itself may slow learning rates during SAT, reducing the activity of SST neurons in somatosensory cortex can overcome this effect to facilitate acquisition of a sensory association.

### SST-depression is not maintained over longer training times

Imaging studies of SST axonal boutons in motor cortex indicate that there may be a slowly-developing reduction in boutons that is associated with continued training in a motor task (Chen et al., 2015). Similar to the increase in reach accuracy during motor learning, we also observed that animals’ performance progressively improved during SAT, measured by the difference in licking frequency between stimulus and blank trials (Figure 1). For example, after 1 day of SAT, 66% of mice showed a significant increase in licking between stimulus and blank trials, but after 5 days, all animals showed significantly more licking to stimulus versus blank trials. To determine whether this increase in performance was associated with a progressive reduction in SST output, we assessed the SST-IPSC in Pyr neurons from both superficial and deep layers after 5 days of SAT.

Longer periods of SAT showed that the large difference in SST output onto L2 Pyr neurons after one day of training can renormalize at longer time points. Although SST-IPSCs were heterogeneous across cells, on average, SST input to L2 Pyr neurons showed no significant difference compared to age-matched controls housed in the training environment but without any coupled sensory input (Figure 7A-D; ACC 219.8±27.2pA versus SAT5 202.5±13.8pA).

**Figure 7.**
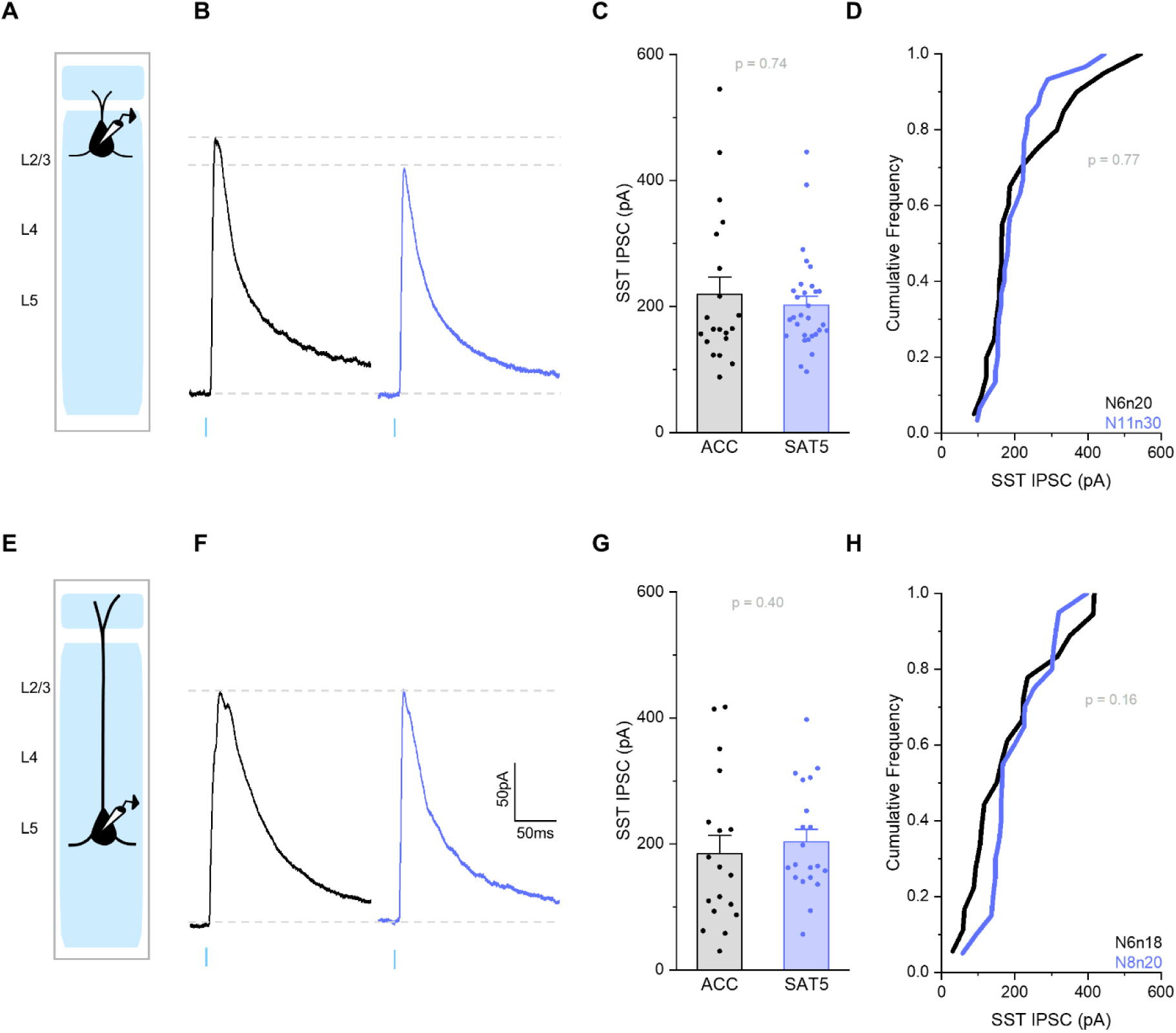
SST inhibition onto L2 Pyr neurons renormalizes after prolonged SAT. (A) Schematic of recording from L2 Pyr neurons. (B) Example ChR2-evoked SST-IPSC (10 sweep average) recorded from a L2 Pyr neuron after 5-6 days of acclimation (ACC; black) and 5 days of SAT (SAT5; blue). (C) Peak amplitude of SST-IPSCs for L2 Pyr neurons from ACC (219.8±27.2pA; N=6 mice, n=20 cells) and SAT5 (202.5±13.8pA; N=11mice, n=30 cells) mice. Bar graphs represent mean+SEM. Mann-Whitney U test (U=283, n=20 and 30, two-tailed). (D) Cumulative distribution histogram of SST-IPSC amplitudes for L2 Pyr neurons from ACC and SAT5 animals. K-S test. (E-H) Same as (A)-(D), but for L5 Pyr neurons (ACC: 185.1±28.5pA; N=6 mice, n=18 cells vs. SAT5: 203.8±19.4pA; N=8 mice, n=20 cells). Mann-Whitney U test (U=151, n=18 and 20, two-tailed). K-S test. *P–*value in figure.

At the same time point (SAT5), we also checked SST output onto L5 Pyr neurons to determine whether inhibition might be modulated at a slower rate here than in superficial layers. However, we did not observe any depression of the SST-IPSC in L5 Pyr neurons even after extended training (Figure 7E-H; ACC 185.1±28.5pA versus SAT5 203.8±19.4pA). These data suggest that changes in the barrel cortex are most pronounced at the early stages of learning and that SST output onto Pyr neurons may be gradually restored with prolonged training regimens.

### SST output depression is correlated with performance in later stages of learning

An advantage of SAT in an automated homecage environment is that large numbers of animals can be trained, and potential correlations between animal behavior and SST output plasticity can be detected. We observed no clear correlation between the amplitude of the SST-IPSC averaged for cells from a single animal and the performance of that animal after 1 day of training (Figure 8A-C). Because analysis of the SST-IPSC was carried out in acute brain slices, we were unable to longitudinally monitor SST output strength within the same animal. However, we were surprised to find after 5 days of training, animal mean SST-IPSC showed a significant, negative correlation with learning, where lower SST-IPSCs were associated with higher performance (Figure 8A, D-E).

**Figure 8.**
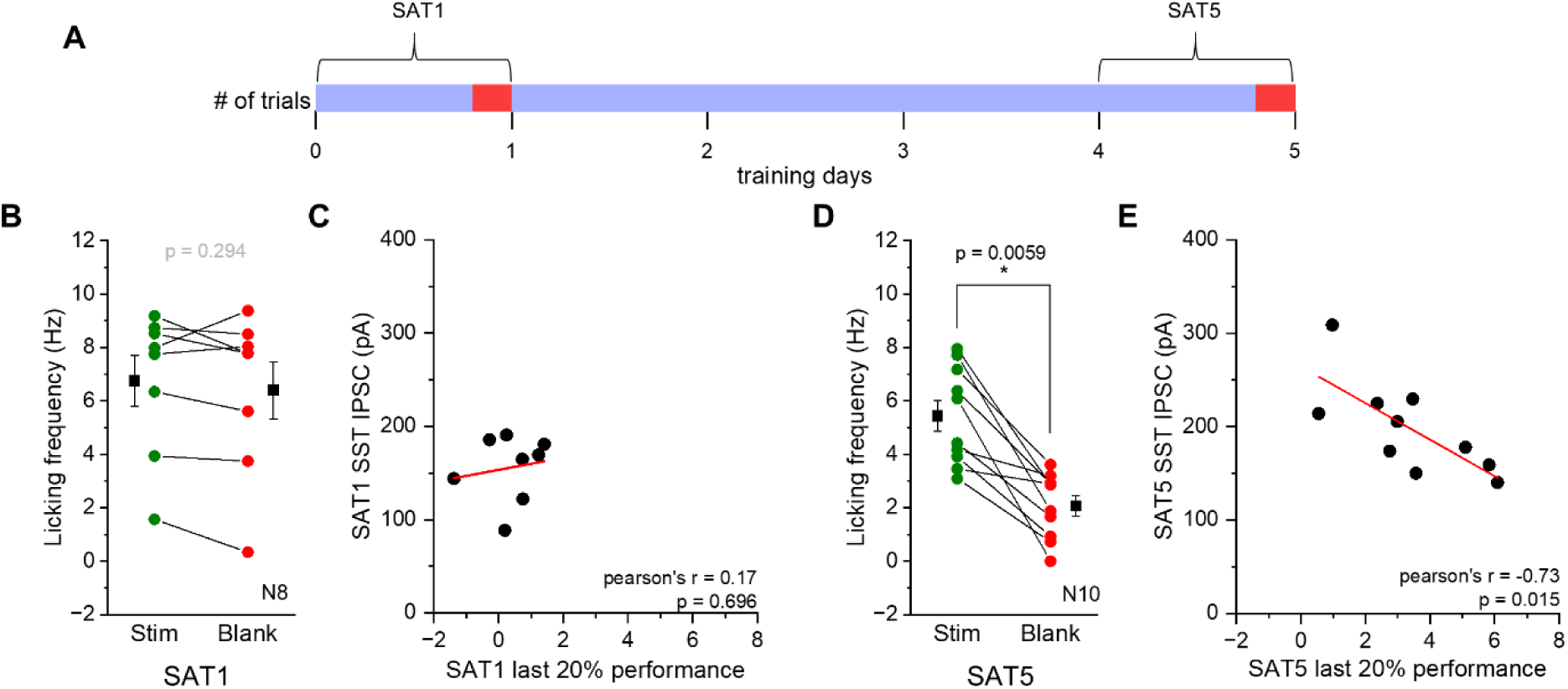
Correlations between SST inhibition and learned behavior. (A) Time windows for behavioral analysis. Average licking frequencies for blank trials and stimulus trials were calculated from the last 20% of total trials (red shade) after 1 day and 5 days of SAT (SAT1 and SAT5, respectively). (B) The comparison of mean anticipatory licking frequency for stimulus (green) and blank (red) trials at the end of 1 day of SAT (SAT1; N=8 mice). Only animals with L2 SST-IPSCs measurements were plotted. Group mean±SEM shown in black symbols (stim: 6.8±1.0Hz vs. blank: 6.4±1.1Hz; N=8 mice). (C) Correlation between performance calculated from last 20% of trials and mean L2 SST-IPSCs for each animal after 1 day of SAT. (D-E) As in (B)-(C), but for animals after 5 days of SAT (stim: 5.4±0.6Hz vs. blank: 2.1±0.4Hz; N=10 mice). Mean anticipatory licking frequency for stimulus and blank trials were compared using a paired-sample Wilcoxon signed rank test. Regression analysis was performed using Pearson correlation. Rho values and two-tailed *p*-value are in figure. \**p<*0.05.

Improvements in behavioral performance after 5 days of training were mainly related to enhanced suppression of licking in “blank” trials (Figure 8B, D). Mean anticipatory lick frequency in stimulus trials was only modestly reduced from the first to the fifth day of training (Figure 1D SAT1 6.2±0.5Hz versus Figure 8D SAT5 5.4±0.6Hz, p=0.17). However, lick frequency in “blank” trials showed a marked reduction over the same time interval, a difference that was highly significant (SAT1 5.6±0.5Hz versus SAT5 2.1±0.4Hz, p=0.004). These data suggest that behavioral improvements with prolonged training may be related to animals being better able to withhold licking in the absence of the sensory stimulus.

Taken together, these data indicate that a reduction in SST-IPSCs in L2 Pyr neurons is not sufficient to drive association learning at SAT1 but may support behavioral modifications during longer training periods.

## Discussion

### Summary

Here we show that neocortical SST neurons can detect behaviorally-relevant signals in the early stages of association learning, reducing their output to selectively disinhibit Pyr neurons in superficial layers of the neocortex. Although prior studies have suggested that inhibition from SST neurons can be dynamically suppressed during specific attentional states or task performance, using acute brain slices from trained animals, we identify a durable reduction of SST-IPSCs that persists outside of the task context. SST synaptic output was decreased in a target-specific manner, onto Pyr neurons but not PV neurons, and this synaptic plasticity could be phenocopied by chemogenetic suppression of SST activity in untrained animals. Importantly, depression of SST output was sensitive to stimulus-reward contingency during association learning since it was absent when stimulus and rewards were uncoupled. Thus, the plasticity of SST inhibition during learning is rapid, sustained, and highly-selective, activated by behaviorally-relevant conditions and controlled by both the presynaptic activity level and the molecular identity of the postsynaptic cell.

### Learning- and experience-dependent plasticity in cortical inhibition

Disinhibition has been proposed as a powerful mechanism to enable circuit plasticity during learning (Barron, 2021; Letzkus et al., 2015) and general evidence for reduced inhibition has been obtained in both sensory and motor learning (Chen et al., 2015; Froemke et al., 2007; Kida et al., 2016; Makino and Komiyama, 2015; Sarro et al., 2015). Anatomical and functional studies across the neocortex indicate that SST inhibition can be dynamically suppressed by the activation of GABAergic VIP neurons (Fu et al., 2014; Gasselin et al., 2021; Pi et al., 2013), suggesting that disinhibition could be a circuit phenomenon that is primarily state-dependent. For example, *in vivo* optogenetic activation of VIP neurons and suppression of SST neurons can facilitate ocular dominance plasticity, a passive form of sensory plasticity (Fu et al., 2015). In addition, both pharmacological and circuit-based interventions *in vitro* indicate that acute suppression of inhibition (Barth et al., 2016; Jiang et al., 2015; Pfeffer et al., 2013) can facilitate synaptic plasticity at glutamatergic synapses (see for example (Canto-Bustos et al., 2022; Glazewski et al., 1998; Williams and Holtmaat, 2019)). However, although a role of short-acting disinhibition in learning is well-documented, there has been little indication that brief periods of disinhibition during the training epoch are sufficient to drive long-lasting changes in inhibitory synaptic function. We show here that SST neurons are selectively sensitive to conditions that drive learning and that this is manifested by long-lasting and target-specific changes in synaptic output.

It has been proposed that increasing inhibition is critical during excitatory synaptic plasticity in order to maintain some balance of excitation and inhibition (Barron, 2021; Okun and Lampl, 2008; Vogels et al., 2011), but these hypotheses have not articulated a source or a target for this inhibition, nor the timescale at which this balancing plasticity should occur. Prior studies have shown that SAT drives the potentiation of both thalamocortical and intracortical excitatory synapses (Audette et al., 2019; Ray et al., 2023). These new data indicate that at least at the early stages of learning, in this task and for L2 Pyr neurons, the potentiation of excitatory synapses is not accompanied by an increase in SST-mediated inhibition.

Do other inhibitory neurons show a similar reduction in output during learning? Although there is evidence for the regulation of PV inhibition by experience, data showing changes in PV output during learning is weaker. Passive sensory experience reduced PV neuron activity *in vivo* (Kato et al., 2015), and circuit plasticity associated with passive sensory experience is impaired with chemogenetic PV neuron activation, suggesting that suppression of PV activity might be important for some kinds of experience-dependent cortical rewiring (Kaplan et al., 2016). However, changes in PV axonal boutons were not robust during motor learning (Chen et al., 2015), and evidence for changes in PV neuron activity during learning has been mixed (Khan et al., 2018; Makino and Komiyama, 2015). The complex pattern of connectivity between different subtypes of interneurons (Barth et al., 2016; Jiang et al., 2015; Pfeffer et al., 2013) as well as their specific input onto Pyr neurons, has complicated efforts to link changes in the activity of different interneuron subtypes to specific synaptic modifications during learning. Further studies to comprehensively investigate synaptic plasticity in inhibitory networks, including direct inhibition onto neocortical Pyr neurons, will be required to understand how changes in inhibition alter information processing in neocortical circuits.

### Layer and target-specific effects

SST neurons have been implicated in learning in prior studies, where a reduction in both their evoked activity as well as their axonal boutons have been identified (Adler et al., 2019; Chen et al., 2015; Khan et al., 2018; Yang et al., 2022) but see (Cummings et al., 2022; Dobrzanski et al., 2022)). Because the target-specificity of these effects has not been assessed, it has been difficult to determine how these changes might influence cortical information processing. For example, a reduction in SST inhibition onto PV neurons might increase PV activity to enhance inhibitory tone in the neocortex. Our data indicate that SST plasticity is target-specific, occurring onto L2 Pyr neurons but not PV neurons, suggesting that the main effect of reduced SST output might be an overall disinhibition of activity in superficial layers. Because evoked SST activity is typically delayed with respect to stimulus onset (Figure S3 and (Gentet et al., 2012; Yu et al., 2019)), this reduction in SST output may directly prolong sensory-evoked responses. It is also possible that reduced SST output in superficial layers could influence cortical activity outside of the task context (Pluta et al., 2019), a process that might be important in consolidating synaptic changes during learning.

Since we were using an SST-Cre transgenic mouse to drive ChR2 expression in all SST neurons, we were unable to evaluate how the composite IPSC recorded in Pyr and PV neurons might be related to synaptic input from different SST subclasses defined by morphology and gene expression (Muñoz et al., 2017; Oliva et al., 2000; Tasic et al., 2018; Urban-Ciecko and Barth, 2016). Because SST neurons in superficial layers are disproportionately Martinotti cells, it is tempting to speculate that this training-dependent plasticity might be concentrated in this SST subclass. If L2 Pyr and PV neurons share inputs from the same class of SST neurons, our data suggest that presynaptic pattern of SST activity by itself is not sufficient to activate synaptic depression, a finding that is relevant to models for inhibitory synaptic plasticity (McFarlan et al., 2023).

Although L5 contains some Martinotti cells, the molecular composition and response phenotypes of SST neurons in infragranular layers are heterogeneous (Muñoz et al., 2017; Yu et al., 2019). In addition, L5 Martinotti cells may be molecularly distinct from those in L2 (Nigro et al., 2018; Tasic et al., 2018). We note that it remains possible that SST input to the apical dendrites of L5 Pyr neurons was altered during training but could not be detected in our assay due to space-clamp restrictions or mechanical transection of the apical dendrite of L5 Pyr neurons during slice preparation. Future studies will be required to resolve whether SST plasticity induced during association learning is restricted by the activity or molecular properties of the presynaptic cell (i.e., Martinotti cells) or the target cell (i.e., L2 Pyr neurons).

### Requirements for SST-output depression during learning

The structure of the sensory training paradigm used here, where the stimulus and reward were temporally decoupled, was designed to examine the intersection of recurrent activity from the stimulus with delayed reward signals, since prior studies have suggested this is important for linking stimuli to subsequent reward information (Hong et al., 2022; Liu et al., 2015; Shuler and Bear, 2006). The selectivity of SST plasticity during SAT but not pseudotraining is consistent with a model that requires the consistent release of neuromodulators during stimulus-reward coupling for synaptic modifications. For example, acetylcholine has been implicated in encoding reward timing in sensory learning (Chubykin et al., 2013; Guo et al., 2019) and both norepinephrine and serotonin have been implicated in inhibitory plasticity (Hong et al., 2022).

Regulation of SST activity may result from reduced excitatory drive, increased inhibition, or changes in the activity of neuromodulators released during training. Because presynaptic VIP neurons can be activated by reinforcement cues (Pi et al., 2013; Szadai et al., 2022), it is tempting to speculate that repeated VIP-mediated suppression of SST firing at the early stages of training might facilitate the weakening of SST output synapses onto L2 Pyr neurons. Our finding that DREADD-suppression of SST activity was sufficient to decrease the SST-IPSC, even in the absence of stimulus-reward pairing, is consistent with this model. Importantly, because direct modulation of SST activity is sufficient to reduce SST-IPSCs, stimulus-reward coupling (with potential state-dependent, neuromodulator activity) may not be required for IPSC depression. The role of specific neuromodulators in the control of SST output plasticity will be an exciting area for further investigation.

### SST inhibition and task performance

SST-IPSCs in L2 Pyr neurons were rapidly reduced at the onset of training, prior to behavioral change in many animals. Our data indicate that changes in SST output by itself is not sufficient to trigger behavioral change. It is likely that learning involves a cascade of synaptic changes that are initiated when environmental stimuli are informative about behavioral outcomes (Barth and Ray, 2019). Indeed, evidence for altered sensory processing prior to learning has been detected in sensory cortex (Audette et al., 2019; Jurjut et al., 2017).

Interestingly, we detected a significant and negative correlation between the SST-IPSC and learning across animals at longer training times. Why was this correlation significant, while the mean SST-IPSC amplitude at later time points appeared to renormalize to cage-matched controls? This may be attributable to several factors: first, the matched control group for SAT5 mice showed a modestly smaller SST-IPSC; second, animals after 5 days of SAT showed a slightly larger SST-IPSC than after one day of training; and third, the variability in SST-IPSC values across animals increased at the longer training interval. However, the significant correlation of SST-IPSCs with performance at SAT5 in a large cohort of animals (10 mice) suggests that this variability is not simply an experimental artifact related to whole-cell patch clamp but reflects differences in the neural circuits that control behavior.

Although we did not directly monitor the activity of SST neurons during sensory stimulation in this study, we did not find that SST-IPSCs progressively reduced with longer training periods. Indeed, SST-IPSC amplitudes were similar to those from cage-matched control animals in later stages of training. These data suggest that SST output depression is not necessarily sustained but may be an early and permissive step during learning-related circuit plasticity in sensory cortex. Importantly, the strong correlation between the SST-IPSC and behavior at SAT5 was not driven by a few animals with poor performance and high SST-IPSCs. Similarly, the lack of correlation at SAT1 could not be attributed to a few animals with high performance and low SST-IPSCs. We hypothesize that in some animals, the signals that maintain reduced SST output may dissipate with longer training periods.

### SST neurons and neocortical function

How does reduced SST inhibition impact cortical function? In contrast to PV neurons, which provide fast feedforward inhibition in superficial layers of sensory cortex (Jouhanneau et al., 2018; Swadlow, 2003), SST neurons are controlled by both local excitation and also feedback inputs from higher-order cortical areas (Chang et al., 2022; Kinnischtzke et al., 2014). Thus, SST neurons may be engaged in the processing of feedback connections to sensory cortex, and a reduction in SST-IPSCs may boost sensory-evoked responses in the local network through both GABAa and GABAb pathways (Urban-Ciecko et al., 2015). SST neurons have also been implicated in receptive field modifications (Adesnik et al., 2012; Gentet et al., 2012; Kato et al., 2015; Pluta et al., 2017), particularly affecting output transformations to L5 (Pluta et al., 2019). It remains to be determined whether this training-initiated reduction in SST output enhances feedforward sensory input or alternately facilitates the influence of feedback inputs to L1 and how these two processes might be intertwined in the early stages of learning.

Experimental evidence suggests that across sensory neocortex, SST neurons can respond to behavioral variables such as reward and novelty (Lee et al., 2022; Ren et al., 2022). Our data indicate that behaviorally-relevant stimuli not only dynamically alter SST responses but also initiate long-term depression at SST outputs. These findings provide valuable constraints to both *in vitro* and computational models of disinhibition during sensory learning (Chiu et al., 2013; Wilmes and Clopath, 2019). They will stimulate further studies to define the neural circuits that enable specific subsets of SST neurons to detect behaviorally-relevant cues during learning and determine how SST output plasticity might be leveraged during early learning for other sensory and motor functions.

## STAR Methods

### Key resource table

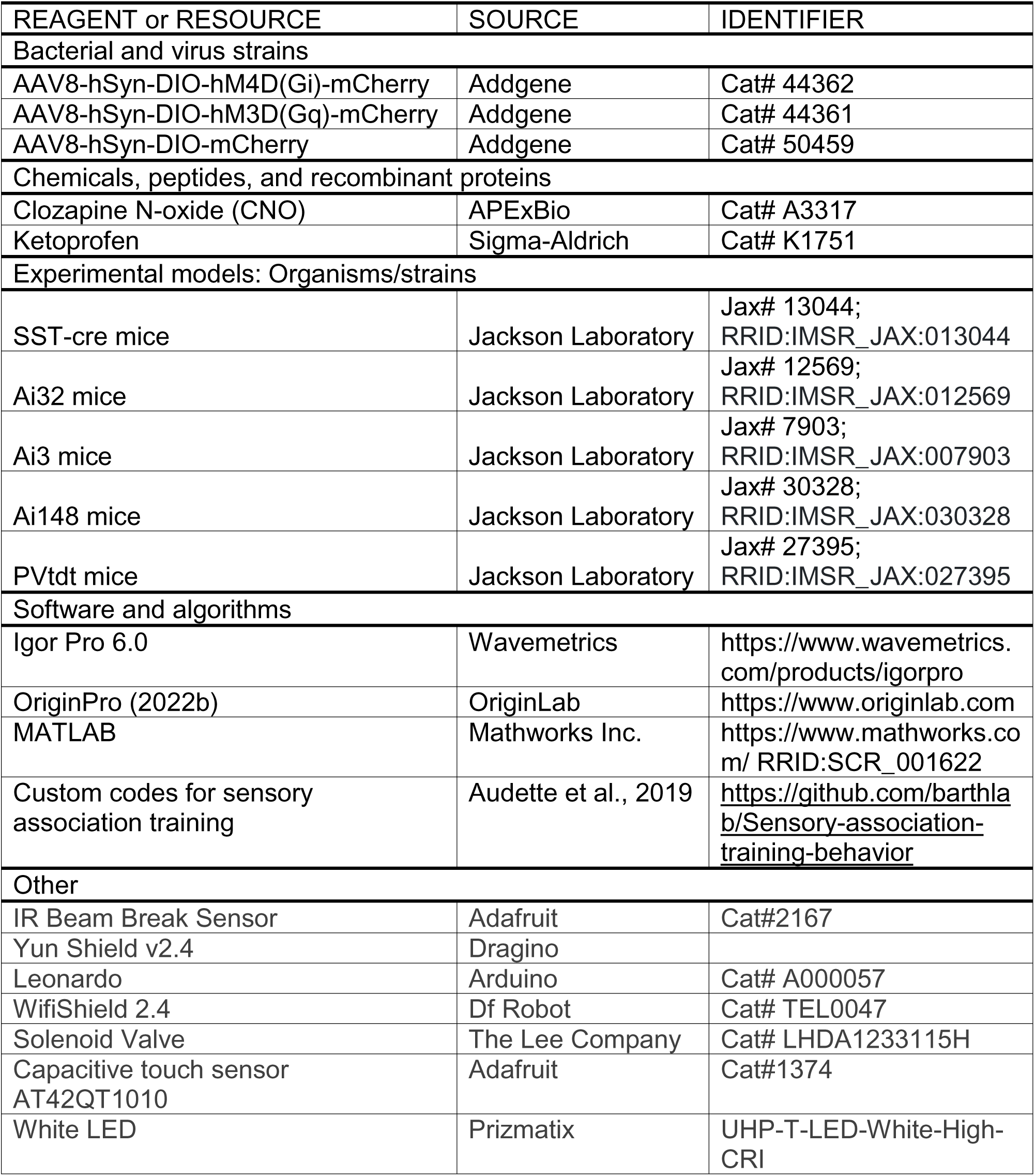

### Animals

All experiments were conducted in accordance with the NIH guidelines and were approved by the Institutional Animal Care and Use Committee at Carnegie Mellon University. SST-IRES-Cre (SST-Cre, Jackson Lab Stock #013044; (Taniguchi et al., 2011)) x Ai32 (Jackson Lab Stock #012569; (Madisen et al., 2012)) mice (male and female, postnatal day (P) 21-31) were used to transgenically express channelrhodopsin-2 (ChR2) in SST neurons. PV-tdTomato (Jackson Lab Stock #027395) x SST-Cre x Ai32 triple-transgenic mice were used to express red fluorescent proteins in PV neurons and ChR2 in SST neurons. SST-Cre x Ai3 (Jackson Lab Stock #007903; (Madisen et al., 2010)) mice were used for a subset of experiments examining the excitability of SST neurons. Juvenile to adult SST-Cre x Ai148 (Jackson #030328; (Madisen et al., 2015)) double-transgenic mice (P50-91) were used for *in vivo* GCaMP6f imaging in SST neurons. Both male and female mice were used.

### Behavior

Animals were single-housed in an automated-home cage for sensory association training (SAT). SAT and pseudotraining were done as previously described (Audette et al., 2019; Bernhard et al., 2020). Animals in all types of training groups were acclimated (ACC) in the training cage for 1-2 days with water delivered in 80% of trials but without any coupled whisker stimulation. Mice that went through fewer than 100 stimulus and reward trials were excluded. Four-hour time bins with fewer than 10 trials were not included in the analysis, since calculations of differences in lick frequency between stimulus (80%) and blank (20%) trials could not be reliably assessed. In some cases, animals were trained, but due to a malfunction of the lick sensor or data upload, the behavioral analysis could not be carried out. Since we could confirm that these animals received the stimulus and also consumed appropriate amounts of water (∼2.5 mL/day), SST-IPSC values for those animals were still included in the dataset.

For chemogenetic experiments, CNO was dissolved in DMSO (0.01 or 0.02 mg/µL) before being added to drinking water. The final concentration of diluted DMSO in the drinking water was ≤0.05%. Mice were weighed before entering the training cage (8-16 g, P22-29). The concentration of CNO (APExBio #A3317) in the drinking water was adjusted based on the mice’s weight and the average volume of water consumption of mice in our study in order to deliver 1 mg/kg/day. Animals were acclimated in the training cage with CNO for 1-2 days before the training started.

### Cranial window surgery

Surgery was done under isoflurane anesthesia (4% for induction, 1.5-2% for maintenance). The mouse was placed on a heating pad with a temperature control system (FHC #40-90-8D) throughout the surgery to maintain body temperature. Dexamethasone (2mg/kg) was injected subcutaneously right before surgery to reduce brain swelling and/or inflammation. Eyes were covered with Puralube Vet Ointment to protect them from drying. The hair on the head was removed with Nair, and the skin was cleaned with povidone and then resected to expose the skull. The skull was scraped with a dental blade (Salvin #6900) to remove the periosteum and to roughen the surface for better attachment of glue. A 3mm diameter circle centered around the left barrel cortex stereotaxic coordinates (bregma (mm): x=−3.5, y=−1.2) was marked with a pen. A thin layer of cyanoacrylate glue (Krazyglue) was applied to the skull, and a custom-made head bracket was attached to the right hemisphere with cyanoacrylate glue and dental cement (Lang Dental #1223PNK). The skull was thinned along the 3mm diameter circle with a dental drill (Dentsply Sirona #780044) to create a craniotomy. Any small bleeding was stopped with saline-soaked gel foam (Pfizer #00009032301). A glass window comprised of a 3mm diameter glass (Warner Instruments #64-0726) attached to a 4mm diameter glass (Warner Instruments #64-0724) by UV adhesive (Norland #717106) was put onto the craniotomy. The window was sealed with 3M Vetbond and Krazyglue. All exposed skull area except for the window was covered with dental cement. A well surrounding the window was built with dental cement to sustain water for the objective lens. Ketoprofen (3mg/kg, Sigma-Aldrich) was injected subcutaneously after surgery. Animals were recovered in a heated cage before returning to their home cage.

### Viral injection

AAV8-hsyn-DIO-hM4D(Gi)-mCherry (Addgene #44362), AAV8-hsyn-DIO-hM3D(Gq)-mCherry (Addgene #44361), or AAV8-hsyn-DIO-mCherry (Addgene #50459) (volume: 0.083-0.4µL) were injected in the barrel cortex (lambda (mm): x= −1.75, y=−1.6, z=−0.3∼−0.5) of SST-Cre or SST-Cre x Ai32 neonates (P1-2). Neonates were anesthetized by 10-12 minutes-long incubation on ice. Two out of 14 animals in Fig 6 were injected with AAV8-hsyn-DIO-mCherry at age P15-18. Mice were anesthetized by isofluorane, and a small craniotomy was created to inject the virus using Nanoject II (Drummond Scientific Company; Broomall, PA). Mice were treated with ketoprofen after injection (5 mg/kg). Virus transduction was checked *post hoc* to verify expression in barrel cortex, and animals with weak hM4D(Gi) expression across the cortical column were excluded from further analysis.

For *in vivo* validation of CNO effects on SST activity, stereotaxic viral injections were performed during cranial window surgery on SST-Cre x Ai148 double transgenic mice (P32-58). A total of 0.44µL pAAV8-hSyn-DIO-hM4D(Gi)-mCherry or AAV8-hsyn-DIO-hM3D(Gq)-mCherry was injected into three different sites in the cranial window around the barrel cortex stereotaxic coordinates (bregma (mm): x=−3.5, y=−1.2, z=−0.3; 18.4nL x 8 times per site). Imaging of SST activity commenced at least 2 weeks after the virus injection.

### Slice preparation

Animals were anesthetized with isoflurane briefly and decapitated between 11 am and 2 pm. 350μm thick off-coronal slices (one cut, 45° rostro-lateral) were prepared in ice-cold artificial cerebrospinal fluid (ACSF) composed of (mM): 119 NaCl, 2.5 KCl, 1 NaH_2_PO_4_, 26.2 NaHCO_3_, 11 glucose, 1.3 MgSO_4_, and 2.5 CaCl_2_ equilibrated with 95%/5% O_2_/CO_2_. Tissues were recovered at room temperature in cutting ACSF for 45 minutes to 1 hour before recording.

### General Electrophysiology

Cortical Pyr neurons were targeted using an Olympus light microscope (BX51WI) and borosilicate glass electrodes (4-9 MΩ pipette resistance) filled with internal solution composed of (in mM): 125 potassium gluconate, 10 HEPES, 2 KCl, 0.5 EGTA, 4 Mg-ATP, 0.3 Na-GTP, and trace amounts of AlexaFluor 594 or 568 (pH 7.25-7.30, 290 mOsm) for morphological confirmation of cell identity. Electrophysiological data were acquired using Multiclamp 700B amplifier (Axon Instruments) and digitized with a National Instruments acquisition interface (National Instruments). Multiclamp and IgorPro 6.0 software (Wavemetrics) with 3kHz filtering and 10kHz digitization were used to collect data. A subset of data collected using 10kHz filtering was still included in the dataset because the amplitude of SST-evoked IPSCs was not affected by the frequency of the filtering threshold. L2 (depth <325µm from the pial surface) and L5 Pyr neurons were targeted based on Pyr morphology with clear apical dendrites in correct laminar orientation.

After breaking into the cells, cells were voltage-clamped at −70mV for 3-5 minutes until the baseline membrane potential stabilized. Pyr cell identity was confirmed by morphology, membrane potential (≤−50mV), input resistance (50-400 MΩ), and regular spiking action potential waveform recorded at current clamp in response to sequential depolarizing current steps (50-700pA, Δ50pA, 500ms duration). ChR2-expressing SST neurons were stimulated with a single light pulse (470nm, 0.58±0.1mW LED, 5ms). The reversal potential of chloride was experimentally determined to be ∼−80mV and was confirmed in all cells included in the analysis. For a subset of experiments, SST-IPSCs were recorded blinded to the experimental conditions. SST-IPSCs were evoked at 0.1

Hz and recorded in voltage clamp at −50mV. Consecutive sweeps (5-10) of SST-evoked IPSCs were averaged, and the average amplitude of SST-evoked IPSCs was measured as the difference between the pre-stimulus baseline and the first peak in the IPSC. Because SAT did not change the excitability of L2/3 SST neurons, the reduction of SST-IPSCs we observed after training is not due to the reduced recruitment of SST neurons during the stimulation period. Indeed, in all cells examined, targeted recording from SST neurons during light illumination was sufficient to drive spiking.

For analysis of SST inputs onto PV neurons, L2/3 PV neurons were targeted using tdTomato reporter fluorescence in PVtdt x SST-Cre x Ai32 triple-transgenic mice. The identity of PV neurons was confirmed by fast-spiking firing phenotype and reporter fluorescence. For a subset of experiments, fast-spiking neurons collected from SST-Cre x Ai32 double-transgenic mice were included as putative PV neurons if optogenetic stimulation did not evoke spikes.

PVtdt x SST-Cre x Ai32, SST-Cre x Ai32, and SST-Cre x Ai3 animals were used to characterize the excitability of L2/3 SST neurons. ChR2- or eYFP-expressing SST neurons were targeted using reporter fluorescence. After breaking into the cell, SST neurons were voltage-clamped at −70mV for 3-5 minutes until the baseline membrane potential stabilized. Action potentials were evoked with depolarizing current steps (25, 50, 100, 150, 200, 250, 300pA, for 500ms duration, 3 sweeps each at 0.1Hz) in current clamp at the cells’ resting membrane potential. Cell identity was confirmed by both YFP fluorescence and low-threshold spiking firing phenotype. Only SST neurons with membrane potential ≤−45mV and a stable membrane potential baseline were included in our analysis. To calculate spiking frequency, depolarizations that exceed 0mV were counted as action potentials. The number of evoked action potentials was calculated by averaging 3 sweeps for each current injection amplitude. To characterize the effect of CNO on the excitability of hM4D(Gi) or hM3D(Gq)-expressing SST neurons, brain slices were washed with 1µM, 5µM, and 10µM of CNO for 5-10 minutes before evaluating evoked firing. Exposure to CNO *in vivo* did not significantly alter intrinsic properties in L2 Pyr neurons as assessed in acute brain slices prepared from these animals.

### Awake head-fixed calcium imaging

We used a 2-photon microscope setup by Femtonics (Femto2D Galvo), equipped with Mai Tai laser MTEV HP 1040S (Spectra-Physics), a 4x air objective lens (Olympus UPLFLN 4X NA 0.13), and a 40x water objective lens (Olympus LUMPLFLN 40XW NA 0.8). Images were acquired with MES software v.6.1.4306 (Femtonics). The mouse was briefly (∼20s) anesthetized with isoflurane (4%) to allow for head fixation under the microscope. The mouse was positioned on top of a wheel to allow for free running during imaging. The mouse was acclimated to the head-fixed position for at least one day, where no stimulus was presented. Blood vessel morphology in 4x brightfield was used to locate the same imaging spot across the imaging period. The pial surface (z=0) was defined as the plane right below the dura matter, which looked like a textured membrane in 40x brightfield. In 40x 2-photon mode, the X, Y, and Z positions of the neurons were aligned to match the previous session image. A 950nm excitation was used to image GCaMP6f signals, and emission fluorescence was detected with green PMT. Laser power and PMT voltage were kept constant within an animal across imaging sessions. Images were acquired at a 5.11 Hz sampling rate with ∼200µm x 250µm field of view and 0.7µm/pixel resolution. The imaging depth was ∼180 - 250µm below pia (L2/3), and two imaging fields per mouse were obtained.

For each day, ∼3-5 minutes after head fixation, two 10-minute imaging sessions with a 1-minute break in between were performed. For each session, either an air puff (500ms, 6psi) or blank (only solenoid click noise) stimulus was delivered using an Arduino interface every 20s (0.05Hz) to the contralateral whiskers. The air puff and blank stimuli had an equal probability of occurring and were randomly interleaved. Typically, ∼10 airpuff stimuli were delivered during each 10-minute session.

After all imaging sessions, the head bracket and window were removed, and the imaging site (confirmed by the morphology of blood vessels) was marked using a glass micropipette containing methylene blue dye. The brain was fixed in 4% paraformaldehyde, flattened, and sectioned tangentially to confirm that the imaging site was in the barrel cortex.

### Image analysis

Images were acquired with MES software v.6.1.4306 (Femtonics). The imaging stack containing all imaging sessions (∼96000 frames) was aligned and segmented with Suite2P (Pachitariu et al., 2017). ROIs were manually selected based on morphology and fluorescence traces calculated by Suite2p (Pachitariu et al., 2017). Any frame that shifted more than 20 pixels in either x or y direction was considered a shifted frame. One or two continuously shifted frames were interpolated with the average value of the previous and the next unshifted frames. If more than two consecutive frames were shifted within a single trial (stimulus onset time±10s), the trial was removed from the further analysis.

Raw fluorescence was extracted for each segmented ROI, and fluorescence signals were neuropil-corrected (F_corrected_=F_ROI_ – 0.7*F_neuropil_). Baseline fluorescence (F_0_) was calculated by averaging the neuropil-corrected signal (F_corrected_) within a 1s time window preceding the stimulus onset of individual trials. The change in fluorescence relative to baseline was calculated as ΔF/F = (F_corrected_ - F_0_)/ F_0_ for every single trial. The stimulus-evoked activity of each neuron was averaged across all stimulus trials within each time window before and after CNO injection.

### Statistics

All statistical tests were performed using OriginPro (Version 2022, Northampton, MA). Anticipatory licking frequencies for responses in stimulus or blank trials and performance (mean±SEM) were averaged in 4-hour time bins to assess animals’ behavior over time. Animals’ performance at the end of a given training day was assessed by calculating anticipatory licking frequency for stimulus and for blank trials for the last 20% of training trials. This took into account variation in the distribution of trials that freely-moving animals carried out at the end of the training period when animals showed variability in trial number due to sleep or episodes of inactivity. Lick frequency for stimulus and blank trials was statistically assessed using a paired-sample Wilcoxon signed rank test. SST-evoked IPSC amplitudes and input resistance are represented as mean±SEM in the graphs. Differences in mean anticipatory licking frequency for stimulus or blank trials between all SAT1 animals (N=29 mice) and SAT 5 animals (N=10 mice) were compared using the Mann-Whitney test.

The average amplitude of SST-IPSCs for the control and experimental group were compared using the Mann-Whitney U test and the Kolmogorov-Smirnov (K-S) test. Membrane potential and rheobase are represented as mean±SD. The effect of experimental conditions on membrane potentials, input resistance, and rheobase of SST neurons was evaluated using the Mann-Whitney U test. The average amplitude of SST-IPSCs in mCherry-injected ACC, hM4D(Gi)-injected ACC, and hM4D(Gi)-injected SAT animals were compared using a one-way ANOVA test. The excitability of SST neurons was compared using two-way repeated measures ANOVA test. Stimulus-evoked SST neuronal activity before and after CNO administration *in vivo* was compared using one-way repeated measures ANOVA test. A *post hoc* Tukey test was used for multiple comparisons following every ANOVA test. The correlation of L2/3 SST IPSCs and the last 20% performance was evaluated using Pearson’s correlation.

---

Table S1 (related to all figures): Numerical values used for figures. Sheet titles correspond to the number of figures.

Table S2 (related to all figures): p-values for statistical tests. Sheet titles correspond to the number of figures.

## Acknowledgments

Special thanks to Joanne Steinmiller for expert animal care, Fernando Bolio for assistance with animal surgeries and histology, and members of the Barth Lab for helpful comments on the manuscript.

## Author contributions

All *in vitro* slice recording experiments were performed by E.P. and D.K. with the help from J.A.C. All chemogenetic experiments were performed by E.P. *In vivo* calcium imaging experiments were performed by M.Z. All data analysis were done by E.P. under the supervision of A.L.B. The manuscript was written by E.P. and A.L.B.

---

**Figure S1.**
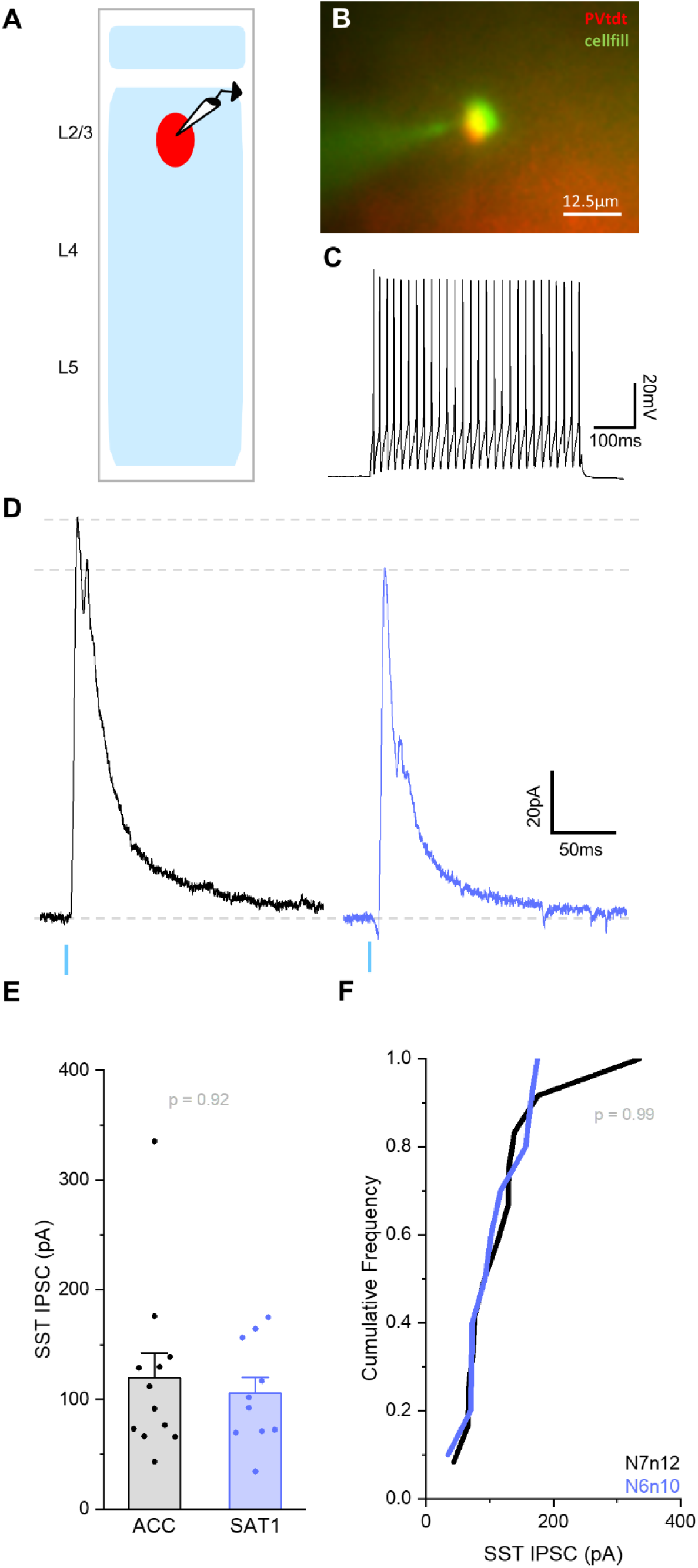
SST inhibition on L2/3 PV neurons is not altered by SAT. (A) Schematic of experimental setup. (B) An example image of targeted L2/3 PV neuron recording. Tdtomato-expressing PV neurons (red) were targeted using a pipette filled with an internal solution containing Alexa 488 (green). (C) The identity of targeted PV neurons was confirmed by their fast-spiking firing phenotype. An example firing response of a L2/3 PV neuron following 500ms duration of 150pA current injection at current clamp. (D) Example ChR2-evoked SST-IPSC (10 sweep average) recorded for a L2/3 PV neuron after 1-2 days of acclimation (ACC; black) and 1 day of SAT (SAT1; blue). (D) Peak amplitude of SST-IPSCs for L2/3 PV neurons from ACC (119.9±22.5pA; N=7 mice, n=12 cells) and SAT1 (105.5±14.8pA; N=6 mice, n=10 cells) mice. Bar graphs represent mean+SEM. Mann-Whitney U test (U=62, n=12 and 10, two-tailed). (E) Cumulative distribution histogram of SST-IPSC amplitudes for L2/3 PV neurons from ACC and SAT1 mice. K-S test. *P–*value in figure.

**Figure S2.**
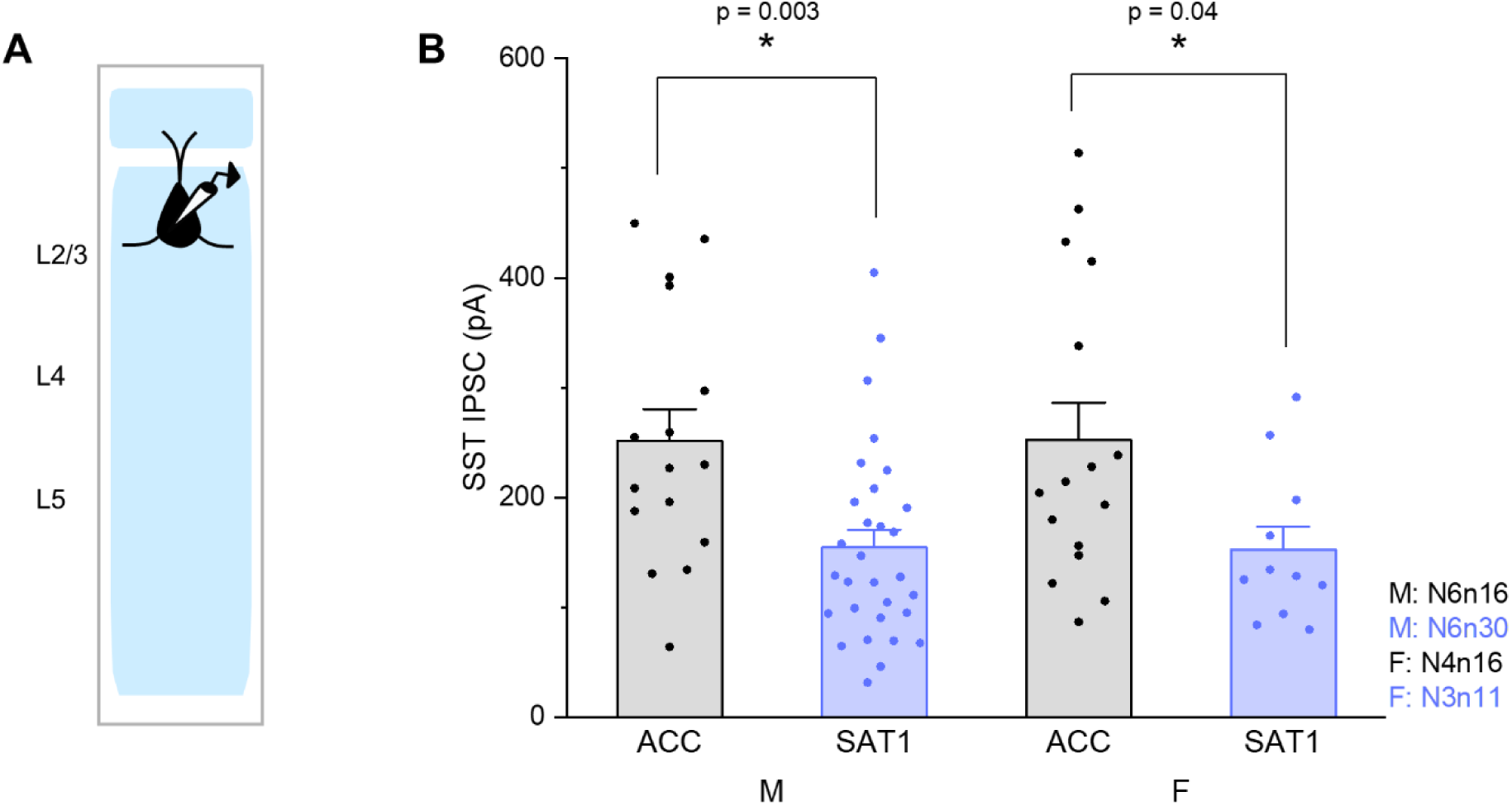
Reduction of L2 SST output is not sex-dependent. (A) Schematic of experimental setup. (B) Mean peak amplitude of SST-IPSCs from L2 Pyr neurons from ACC and SAT1 mice, divided into males (ACC: 251.8±28.8pA; N=6 mice, n=16 cells vs. SAT1: 154.6±16.2pA; N=6 mice, n=30 cells) and females (ACC: 252.6±34.1pA; N=4 mice, n=16 cells vs. SAT1: 152.6±21.0pA; N=3 mice, n=11 cells). Bar graphs represent mean+SEM. Mann-Whitney U test between ACC and SAT1 within the same sex. (Males: U=369, n=16 and 30, two-tailed; Females: U=130, n=16 and 11, two-tailed). *P–*value in figure. \**p*<0.05.

**Figure S3.**
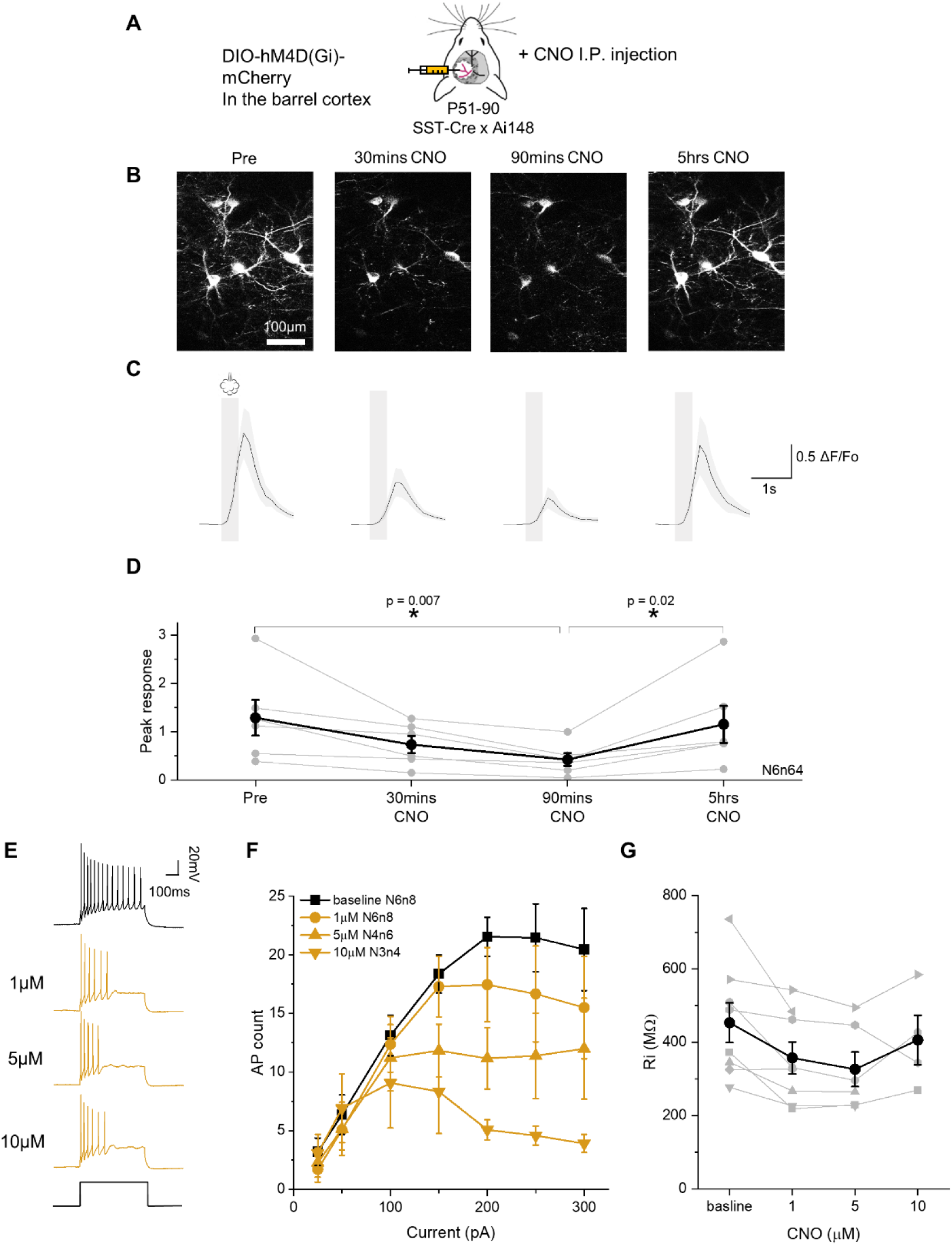
*In vivo* CNO reduces stimulus-evoked activity in hM4D(Gi)-expressing SST neurons. (A) Schematic of the experimental setup. DIO-hM4D(Gi)-injected SST-Cre x Ai148 mice were used. Intraperitoneal (*I.P.*) injection was performed before *in vivo* imaging to deliver 1mg/kg/day CNO. (B) Example field of view. Changes of calcium signals in hM4D(Gi)-expressing SST neurons in response to air puff delivery were monitored before (pre), after 30 minutes, 90 minutes, and 5 hours of CNO *I.P.* injection. (C) Population average of stimulus-onset-aligned ΔF/F traces for the same group of SST neurons. Grey bars mark the duration of airpuff. N=6 mice, n=64 cells. (D) Changes in the averaged peak responses of all hM4D(Gi)-expressing SST neurons before and after CNO *I.P.* injection. Grey lines represent the averaged peak responses of individual mice. A black line (mean±SEM) represents average peak responses from all mice. One-way repeated measures ANOVA. F_(1,5)_=12.5, *p*-value=0.02. Tukey’s *p*-value<0.05 in figure. \**p*<0.05. (E) Example of 150pA current injection -evoked firing from L2/3 hM4D(Gi)-expressing SST neurons in response to 1, 5, and 10µM of CNO wash in acute brain slices. (F) F-I curve of L2/3 SST neurons before (black) and after (yellow) CNO application. Mean±SEM. (G) Input resistance change of L2/3 SST neurons in response to different CNO concentrations. Grey lines represent individual cells, and the black line represents mean±SEM.

**Figure S4.**
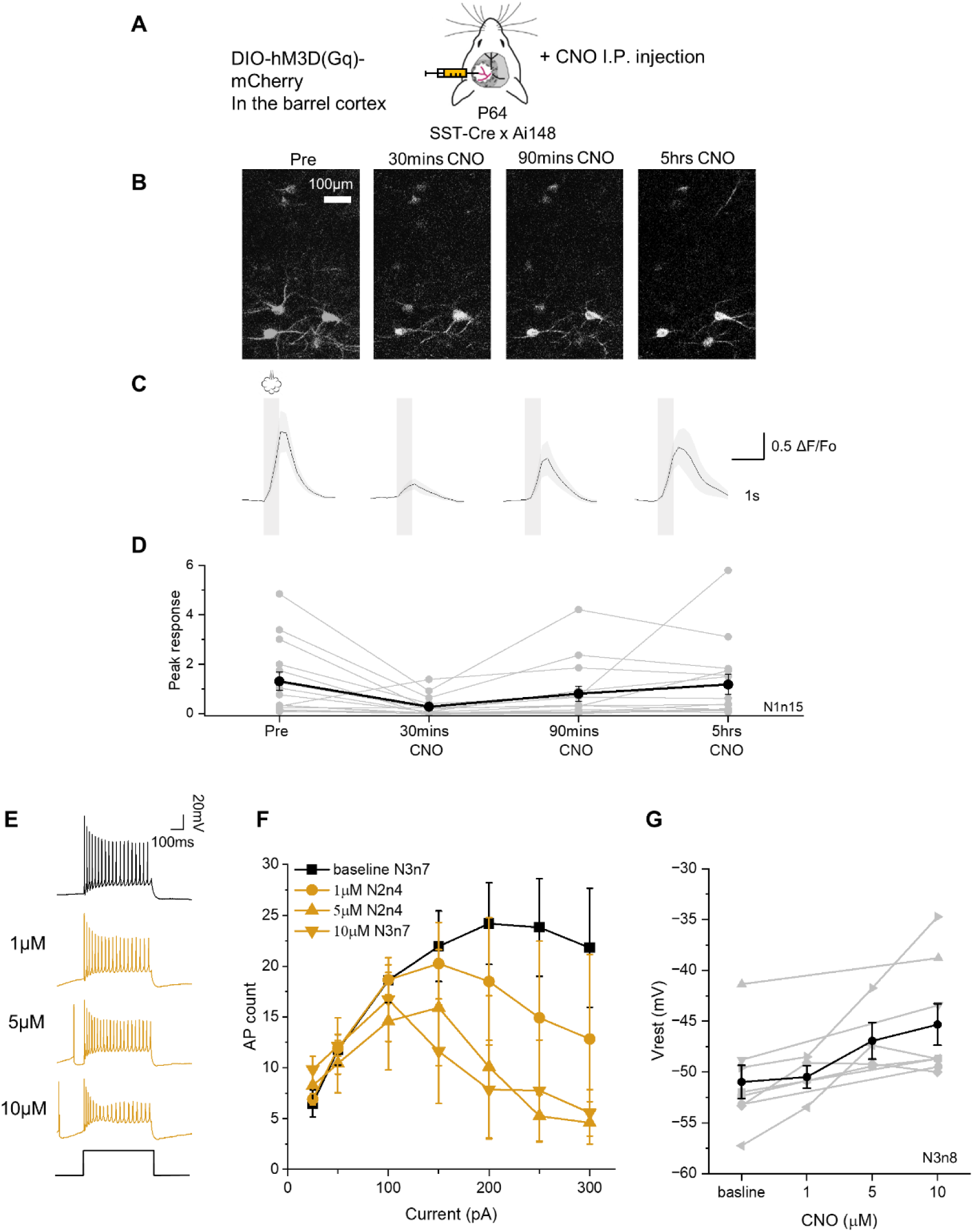
*In vivo* CNO reduces stimulus-evoked activity in hM3D(Gq)-expressing SST neurons. (A) Schematic of the experimental setup. DIO-hM3D(Gq)-injected SST-Cre x Ai148 mice were used. Intraperitoneal (*I.P.*) injection was performed before *in vivo* imaging to deliver 1mg/kg/day CNO. (B) Example field of view. Changes of calcium signals in hM3D(Gq)-expressing SST neurons in response to air puff delivery were monitored before (pre), after 30 minutes, 90 minutes, and 5 hours of CNO *I.P.* injection. (C) Population average of stimulus-onset-aligned ΔF/F traces for the same group of SST neurons. Grey bars mark the duration of airpuff. N=1 mouse, n=15 cells. (D) Changes in the averaged peak responses of all hM3D(Gq)-expressing SST neurons before and after CNO *I.P.* injection. Grey lines represent the averaged peak responses of individual cell. A black line (mean±SEM) represents average peak responses from all cells. (E) Example of 150pA current injection-evoked firing from L2/3 hM3D(Gq)-expressing SST neurons in response to 1, 5, and 10µM of CNO wash in acute brain slices. (F) F-I curve of L2/3 SST neurons before (black) and after (yellow) CNO application. Mean±SEM. (G) Membrane potential change of L2/3 SST neurons in response to different CNO concentrations. Grey lines represent individual cells, and the black line represents mean±SEM.

**Table S3(related to all figures):**
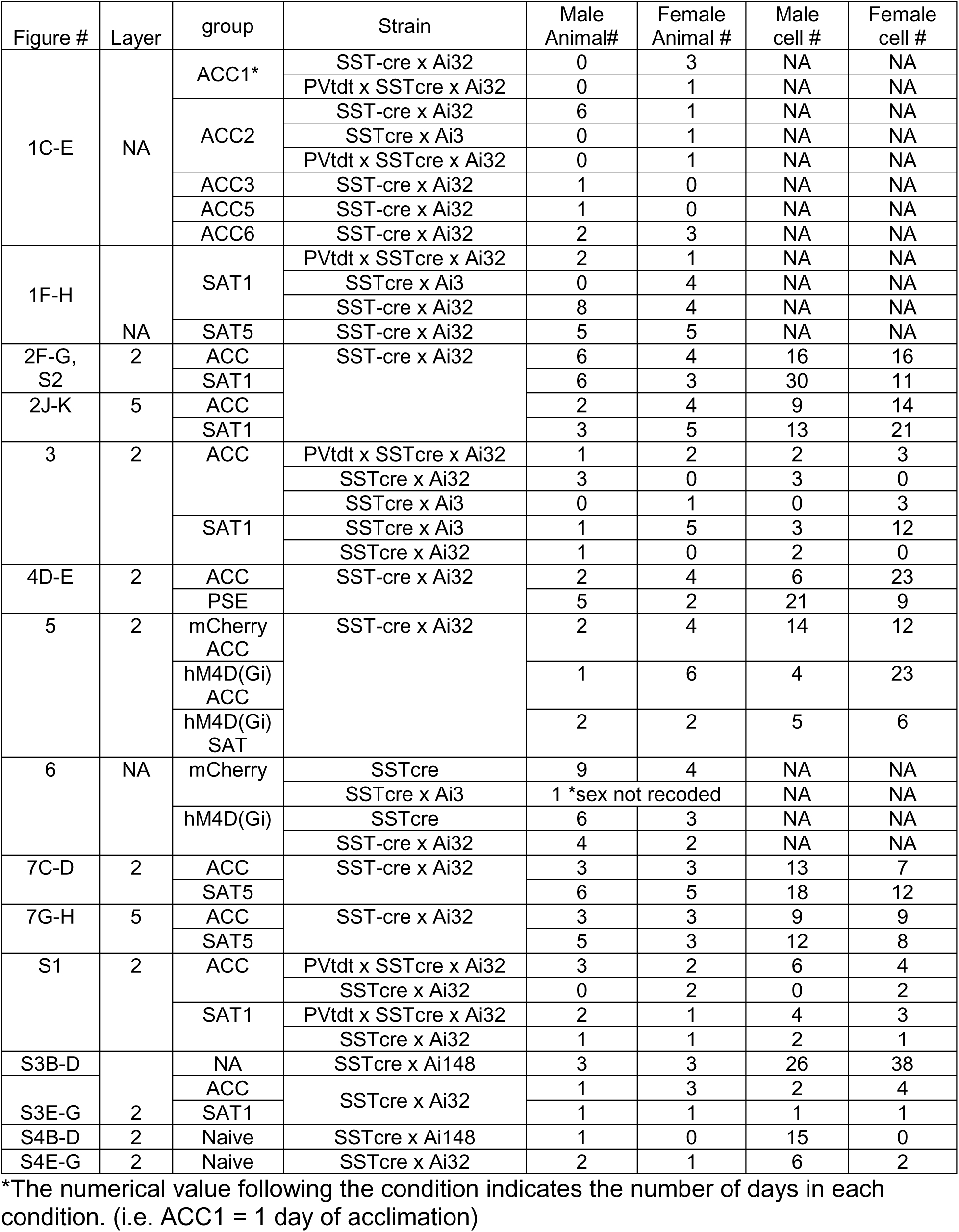
Summary of the animals’ sex for the experiments done for each figure.

